# The Discovery and Characterization of 1,4-Dihydroxy-2-naphthoic Acid Prenyltransferase Involved in the Biosynthesis of Anthraquinones in *Rubia cordifolia*

**DOI:** 10.1101/2023.09.13.557651

**Authors:** Changzheng Liu, Ruishan Wang, Sheng Wang, Tong Chen, Chaogeng Lyu, Chuanzhi Kang, Xiufu Wan, Juan Guo, Luqi Huang, Lanping Guo

## Abstract

Anthraquinones constitute the largest group of natural quinones, which are used as safe natural dyes and have many pharmaceutical applications. In plants, anthraquinones are biosynthesized through two main routes: the polyketide pathway and the shikimate pathway. The shikimate pathway primarily forms alizarin-type anthraquinones, and the prenylation of 1,4-dihydroxy-2-naphthoic acid is the first pathway-specific step. However, the prenyltransferase responsible for this key step remains uncharacterized. In this study, the cell suspension culture of *Rubia cordifolia*, a plant rich in alizarin-type anthraquinones, was used in target prenyltransferase mining. The microsomal protein prepared from the cell suspension culture prenylated 1,4-dihydroxy-2-naphthoic acid to form 2-carboxyl-3-prenyl-1,4-naphthoquinone and 3-prenyl-1,4-naphthoquinone. Then a candidate gene belonging to UbiA superfamily, *RcDT1*, was discovered to account for the prenylation activity. Substrate specificity studies revealed that the recombinant RcDT1 recognized naphthoic acids primarily, followed by 4-hydroxyl benzoic acids. The prenylation activities of *R. cordifolia* microsomes and the recombinant RcDT1 were both strongly inhibited by 1,2- and 1,4-dihydroxynaphthalene. The plastid localization and root-specific expression further confirmed the participation of *RcDT1* in anthraquinone biosynthesis. The phylogenetic analyses of *RcDT1* and its rubiaceous homologs indicated that DHNA-prenylation activity evolved convergently in Rubiaceae via recruitment from the ubiquinone biosynthetic pathway. The discovery and evolutionary studies of *RcDT1* provide useful guidance for identifying additional and evolutionarily varied prenyltransferases which enable entry into quinones derived from shikimate pathway. Moreover, these findings will have profound implications for understanding the biosynthetic process of the anthraquinone/naphthoquinone ring derived from shikimate pathway.

## Introduction

Natural anthraquinones (AQs) and their derivatives are the largest group of natural quinones, most of which are produced by plants, lichens and fungi (Diaz-Muñoz et al. 2018; Wang et al. 2023a). Plant-derived AQs have the most common structure of 9,10-AQ with three rings and can be classified as emodin-type AQs or alizarin-type AQs based on the presence of two or one hydroxylated ring in the structure (Wang et al. 2023a). Plant-derived AQs are mainly distributed in the families Polygonaceae, Rubiaceae, Leguminosae, Rhamnaceae, Scrophulariaceae, Liliaceae, Verbenaceae and Valerianaceae (Thomson 1991; Wang et al. 2023a). AQ-rich plants in these families, for instance, *Rheum palmatum*, *Polygoni multiflora*, *Rubia cordifolia*, *Morinda officinalis*, and *Cassia obtusifolia* have been used to treat various diseases for centuries in China. Modern pharmacological studies have proved that AQs had potent anti-cancer, anti-pathogenic microorganisms, anti-oxidation, anti-osteoporosis, anti-inflammatory, anti-injury, anti-depression, anti-constipation and other pharmacological activities (Wang et al. 2021a). In addition, due to the inclusion of numerous chromophore groups and auxochrome groups in the basic skeleton, natural AQs are dark in color and have been used as natural dyes since ancient times.

In higher plants, two main distinct biosynthetic pathways leading to AQs have been reported: the polyketide pathway for emodin-type AQs and the shikimate pathway for alizarin-type AQs (Diaz-Muñoz et al. 2018; Wang et al. 2023a). Tracer studies and phytochemical analyses support the view that alizarin-type AQs found in Rubiaceae, Bignoniaceae and Verbenaceae are formed by the latter pathway (Burnett and Thomson 1968a, 1968b; Han et al. 2001). This pathway comprises three parts (Fig. 1): (1) conversion of shikimate to the naphthoquinone (NQ) head-group precursor 1,4-dihydroxy-2-naphthoic acid (DHNA), which gives rise to ring A and B of AQs; (2) synthesis of 3,3-dimethylallyl pyrophosphate (DMAPP), which is carried out via the MEP pathway in Rubiaceae; and (3) attachment of DMAPP to the head group and ring closure to produce the ring C. The same biosynthetic logic also applies to NQs of plants, for example shikonin and chimaphilin (Meyer et al. 2021; Widhalm and Rhodes 2016). In these compounds ring A is derived from shikimate. The prenylation step and subsequent cyclization result in the formation of ring B (Fig. 1). Obviously, the prenylation step is of key importance for the formation of AQ/NQ structure in shikimate pathway.

**Figure 1.**
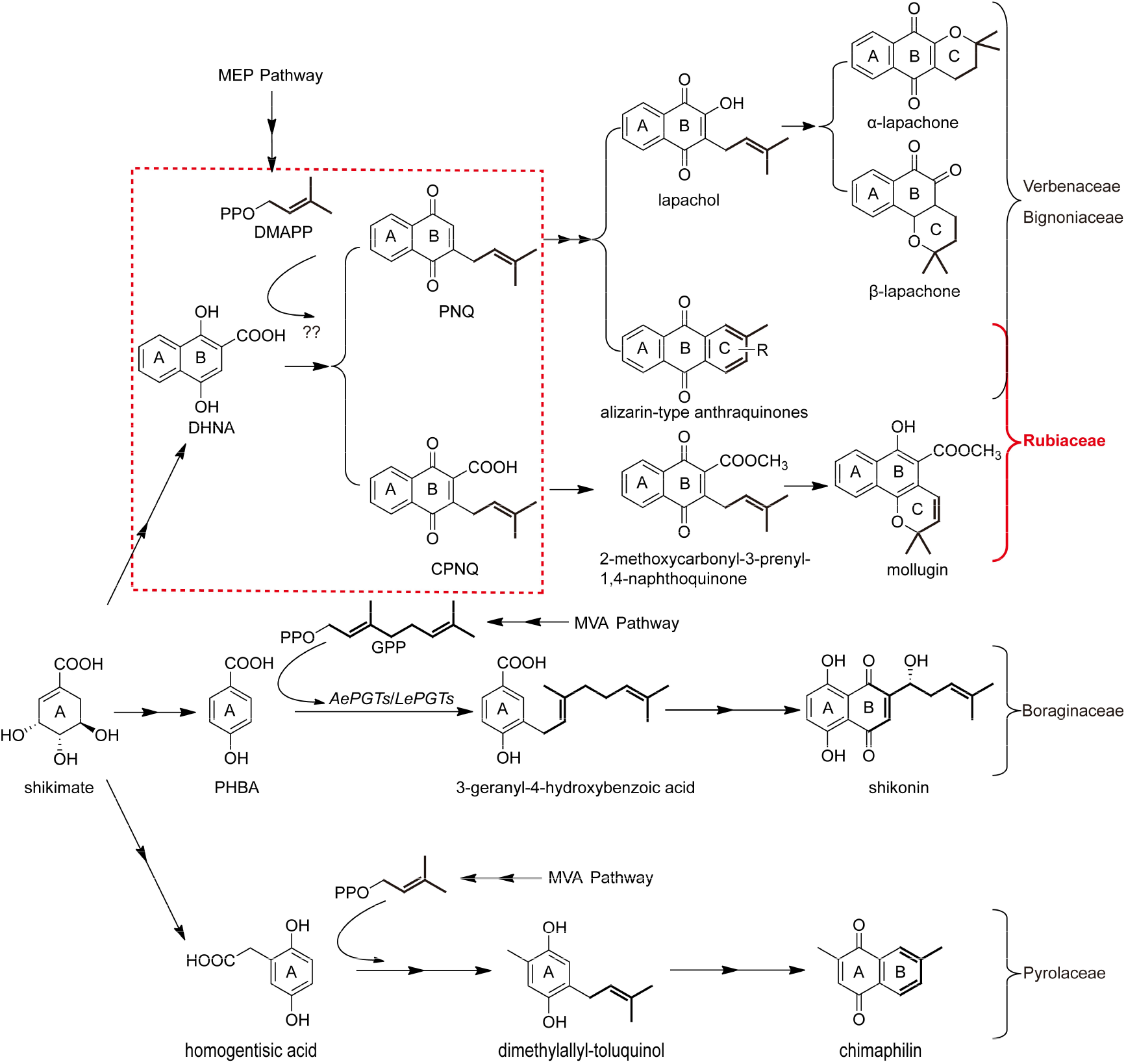
Biosynthetic pathways of shikimate-derived naphthoquinones and anthraquinones in higher plants. The problem addressed in this study is highlighted in red-dotted box. DMAPP, dimethylallyl diphosphate; GPP, geranyl diphosphate; DHNA, 1,4-dihydroxy-2-naphthoic acid; CPNQ, 2-carboxyl-3-prenyl-1,4-naphthoquinone; PNQ, 3-prenyl-1,4-naphthoquinone; PHBA, 4-hydroxybenzoic acid.

The prenylation reaction in plants is generally catalyzed by membrane-bound prenyltransferases (PTs) belonging to the UbiA superfamily (An et al. 2023). Members of this superfamily are widely present in prokaryotes and eukaryotes and produce lipophilic compounds both in the primary metabolism (e.g., ubiquinones and menaquinones) and specialized metabolism (e.g., prenylated flavonoids, coumarin, and stilbenoid) (Li 2016). The first DHNA PT was identified in *Escherichia coli* as a member of UbiA family, i.e. MenA, a DHNA-octaprenyltransferase involved in the biosynthesis of menaquinone (Suvarna et al. 1998). Then DHNA-phytyltransferases, homologs of MenA from cyanobacteria and higher plants, have been confirmed to be involved in the synthesis of phylloquinone (Johnson et al. 2000, 2001; Shimada et al. 2005). In animals, UBIAD1, a homolog of MenA, is the PT for menaquinone-4 (MK4) biosynthesis and associated with many physiological processes and diseases (Nakagawa et al. 2010). However, the gene encoding the DHNA-dimethylallyltransferase (DHNA-DT) that leads to the formation of ring C in alizarin-type AQs is yet to be discovered. Its genetic information, enzymatic characteristics, reaction mechanism, resulting intermediates, and evolution remain enigmatic.

In this context, cell suspension culture of *R. cordifolia*, a representative species of Rubiaceae, was established in our laboratory and used as a material for analyzing the biosynthetic pathway of alizarin-type AQs. The microsomal fraction prepared from this material was found to exhibit DHNA-DT activity. Then the PT gene *RcDT1* was cloned from the cell suspension culture and proved to be responsible for the activity. Unlike MenA-catalyzed prenylation, the prenylation of DHNA by RcDT1 occurred at C-3 position and was accompanied by a spontaneous naphthol to naphthoquinone oxidation and decarboxylation. Functional validation and phylogenetic analysis for the homologs of *RcDT1* in rubiaceous plants *Coffea arabica*, *Coffea canephora* and *M. officinalis* indicated that DHNA-DTs in Rubiaceae were recruited from the ubiquinone biosynthetic pathway and evolved convergently. These results have deepened the understanding of AQ biosynthesis in the shikimate pathway and would have profound implications for elucidating the subsequent cyclization steps in AQ/NQ biosynthesis by the shikimate pathway.

## Results

### Enzymatic activity of *R. cordifolia* microsomes

Plant cell suspension cultures are fundamental research tools for elucidating the biosynthesis of plant specialized metabolites. In order to explore the DHNA-prenylation activity in alizarin-type AQ biosynthesis, *R. cordifolia*, a representative species of Rubiaceae, was chosen for research and its cell suspension culture was established. The microsomal fraction was extracted and incubated with DHNA (**1**), the biosynthetic intermediate of alizarin-type anthraquinone, together with the prenyl donor DMAPP and the cofactor Mg^2+^. As a result, a major product **1a** and a minor product **1b** were detected simultaneously at retention times of 1.8 min and 4.4 min (Fig. 2B). Product formation was strictly dependent on the presence of DHNA, DMAPP, Mg^2+^, and the active microsomal protein. HPLC-UV/ESI MS analysis revealed that the molecular weight of the major product (**1a**) was 66 amu higher than DHNA (**1**), indicating that DHNA (**1**) was prenylated (68-amu increase in molecular weight) and oxidized to naphthoquinone (2-amu decrease in molecular weight) (Fig. 2D and 2E). Previously, tracer studies and phytochemical analyses have confirmed that prenylation during the biosynthesis of alizarin-type AQs in Rubiaceae occurred at the position C-3 of DHNA (**1**) (Heide and Leistner 1981; Inoue et al. 1979). Accordingly, the chemical structure of the major product **1a** was inferred to be 2-carbonyl-3-prenyl-1,4-naphthoquinone (CPNQ), which was likely to be a branch point in the biosynthetic pathway leading to AQ and naphthohydroquinone (NHQ, e.g., mollugin) derivatives (Fig. 1). Through comparing with an authentic standard using HPLC-UV/ESI-MS analysis, the minor product (**1b**) was identified as 3-prenyl-1,4-naphthoquinone (PNQ, i.e. deoxylapachol) (Fig. 2B and 2F). Both the molecular weight of **1b** (44-amu less than **1a**) and the mass spectrometry fragmentation of **1a** and **1b** supported that decarboxylation of **1a** produced **1b** (Fig. 2E and 2F). Since β-keto acid decarboxylation was known in both organic chemistry and biocatalysis, PNQ (**1b**) was assumed to be formed through spontaneous decarboxylation of CPNQ (**1a**) (Wellington et al. 2012). This prenylation reaction mode was different from that observed in MenA, which catalyzed the C-2 prenylation of DHNA (**1**) (Baldwin et al. 1973). Due to the instability of CPNQ (**1a**), the attempt to isolate **1a** in the solid state from the reaction system failed and the decarboxylation of **1a** has not been further studied *in vitro*.

**Figure 2.**
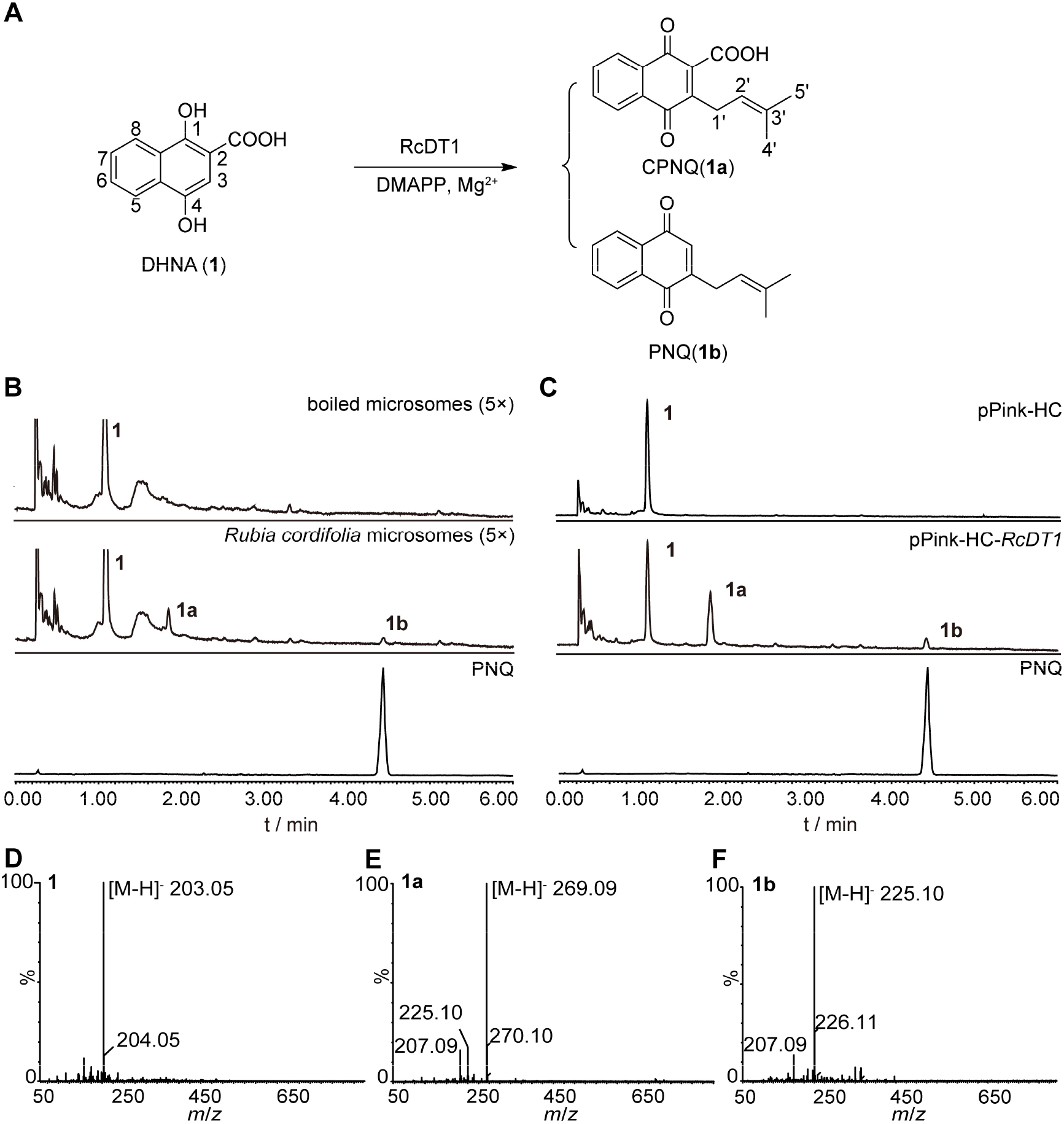
Functional characterization of *R. cordifolia* microsomes and recombinant RcDT1. The detection wavelength is set at 254 nm. PNQ is used as an authentic standard. A, Reaction mode diagram by *R. cordifolia* microsomes. The chemical structures of the compounds and corresponding peaks in UPLC chromatogram are marked as **1**, **1a**, and **1b**. B, UPLC traces for the reactions containing DHNA (**1**) with active or boiled *R. cordifolia* microsomes. C, UPLC traces for the reactions containing DHNA (**1**) with recombinant RcDT1 microsomes or microsomal extract from yeast transformed with empty vector. C, MS spectrum of DHNA (**1**). D, MS spectrum of the product **1a**. E, MS spectrum of the product **1b**.

### Cloning and characterization of *RcDTs* from the cell suspension culture of *R. cordifolia*

To mine the DHNA (**1**) PT gene(s) responsible for the prenylation catalyzed by cultured *R. cordifolia* cells, a RNA-seq library was prepared and then used for transcriptome analysis. *AtABC4* (a DHNA phytyltransferase homologous to MenA in *Arabidopsis thaliana*), *LePGT1* [a 4-hydroxybenzoic acid (PHBA) geranyltransferase (PGT) in *Lithospermum erythrorhizon*], *AePGT6* (a PGT in *Arnebia euchroma*), and *SfN8DT-1* (a naringenin DT in *Sophora flavescens*) were employed as queries to retrieve the transcriptome database independently. As a result, the search yielded six putative aromatic PT genes, *RcDT1 – 6*. No candidate gene with a certain similarity (E-value<1) to *AtABC4* was found.

The encoded polypeptides of *RcDT1 – 6* had five to eight putative transmembrane α-helices as predicted by the DeepTMHMM program. A multiple alignment of the encoded polypeptides and query UbiA enzymes was shown in Fig. S1. The polypeptides possessed two conserved aspartate-rich motifs N(Q/D)*XX*D*XXX*D and D*XX*D*XXX*D. The former motif is commonly thought to be involved in the binding of prenyl diphosphate (Yazaki et al. 2009). The latter motif exhibited a high degree of variability in the alignment. SignalP-6.0 predictions indicated that the polypeptides possess N-terminal signal peptides in the length range of 20-50. The above data matched the typical structural characteristics of plant aromatic PTs (Winkelblech et al. 2015).

Previous studies had shown that N-terminal truncation usually increased the enzymatic activity of heterologously expressed UbiA enzymes. Consequently, *RcDT1 – 6* with an N-terminal truncation of 24 amino acids were expressed in *Pichia pastoris* to detect their prenylation activity against DHNA (**1**). As a result, incubation of DHNA (**1**) and DMAPP with the recombinant *RcDT1* microsomes in the presence of Mg^2+^ led to the detection of CPNQ (**1a**) and PNQ (**1b**) (Fig. 2C). Product formation was strictly dependent on the presence of **1**, DMAPP, Mg^2+^, and the active recombinant *RcDT1* microsomal protein. This was consistent with the results obtained with *R. cordifolia* microsome, and the transformation efficiency was increased significantly (Fig. 2B and 2C). *RcDT1* shared identities of 53.11% and 50.16% at the amino acid level with *AePGT6* and *LePGT1* respectively, suggesting that this gene might have a similar evolutionary origin to boraginaceous PGTs. We further compared the activities of truncated and untruncated *RcDT1*. The activity of truncated *RcDT1* with a 24 or 48 or 72 amino acid deletion from the start codon was not significantly different, which was much higher than that of untruncated *RcDT1* (data not shown). In subsequent experiments, the truncated *RcDT1* with a 72 amino acid deletion at the N-terminal was selected to study the biochemical properties. Due to the lack of activity when using DHNA (**1**) as the substrate, the candidate genes *RcDT2 – 6* were not investigated further.

### Substrate specificities of the recombinant RcDT1

To investigate the prenyl acceptor specificities of recombinant RcDT1, we tested 23 potential acceptors including various types of naphthoic acids and a naphthalene (**1** – **8**), benzoic acids (**9** – **18**), phenylacetic acids (**19**, **20**), and phenylpropanes (**21** – **23**). (Fig. 3B). The results showed that RcDT1 could accept both naphthoic acids **1**-**5** and benzoic acids **9**-**13** as prenyl acceptors (Fig. 3C). The enzymatic products of **2**, **3**, **9**, and **11** were prepared from larger scale assays and subjected to MS and NMR spectroscopic analyses (Fig. 3A). Chemical structure of the enzymatic product of **10** was confirmed by UPLC-UV/ESI MS analysis. The prenylated products of **4**, **5**, and **13** were not prepared successfully due to the relative low yields. For naphthoic acid substrates, one dimethylallyl moiety was introduced into the carboxyl-bearing ring at the position C-3 or C-4 (Fig. 3A). For benzoic acid substrates one dimethylallyl moiety was regioselectively introduced into the *ortho* position of 4-hydroxyl group (Fig. 3A).

**Figure 3.**
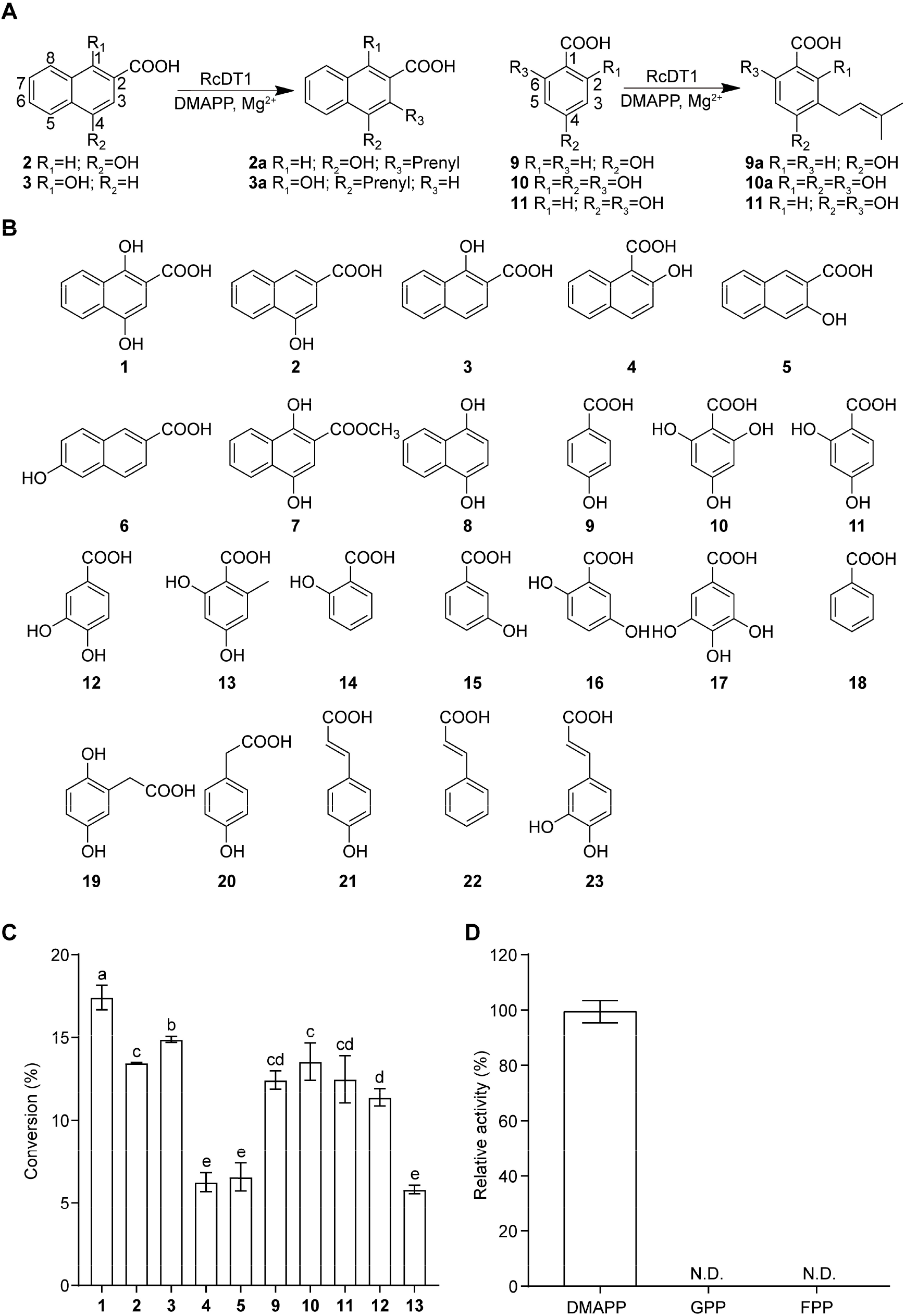
Substrate specificities of the recombinant RcDT1. A, The reactions catalyzed by the recombinant RcDT1. B, Chemical structures of the compounds for the substrate specificity analysis. C, Relative enzymatic activities with the substances in Figure 3B as the prenyl acceptors and DMAPP as the prenyl donor. One-way ANOVA is used to compare the conversion. Different lowercase letters indicate significant differences (*P*<0.05). D, Relative enzymatic activities with DMAPP, GPP and FPP as the prenyl donors and DHNA (**1**) as the prenyl acceptor. N.D., not detected.

The reactivities of RcDT1 toward different naphthoic acids and benzoic acids provided insight into the reaction mechanism and enzyme selectivity. First of all, the relative conversion rate of DHNA (**1**) was the highest among all the substrates, indicating that it was the natural substrate of RcDT1. Regarding the recognition of DHNA (**1**), the existence of unsubstituted 2-carboxyl was crucial, because the absence (**8**) or the methylation (**7**) of 2-carboxyl led to no enzymatic product generation. The prenylation could still take place in C-1/3/4 hydroxylated 2-naphthoic acid (**2**, **3**, and **5**) and 2-hydroxy-1-naphthoic acid (**4**). However, loss of the hydroxyl groups in the carboxyl-bearing ring led to no prenylation (**6**). The unsubstituted ring in the recognized naphthoic acids (**1** – **5**) also played an important role in substrate recognition, because the benzoic acid substructure fragments (**14** – **16**) of **1** – **5** could not be prenylated. To sum up, naphthoic acids with hydroxyl(s) in the carboxyl-bearing ring were putative substrates of RcDT1.

The prenylations are electrophilic substitution reactions and the reaction mechanism includes a prenyl carbocation. As observed in the experiments, the prenyl substitution occurred at the position with sufficient electron density (e.g., C-4 of **3**). It is worth noting that decarboxylation is not observed for the other available naphthoic acids except DHNA (**1**). This result could be interpreted that β-keto in CPNQ (**1a**), which was generated by spontaneous oxidation, was a prerequisite for the decarboxylation. Therefore, the reaction mechanism of DHNA prenylation by RcDT1 could be summarized as follows: dimethylallyl moiety was firstly introduced into the C-3 of DHNA; then the naphthol structure was oxidized to NQ spontaneously to produce CPNQ (**1a**); at last, the decarboxylation of CPNQ (**1a**) led to the generation of PNQ (**1b**).

In addition to naphthoic acids, RcDT1 also exhibited relatively high catalytic activity toward benzoic acids (Fig. 3C). For the recognition of benzoic acid substrates, RcDT1 recognized only the benzoic acids with 4-hydroxyl group (**9**–**13**). And the prenylation occurred exclusively at an adjacent position of the 4-hydroxyl group (Fig. 3A). This prenylation reaction mode of RcDT1 for 4-hydroxyl benzoic acids resembled that of PHBA polyprenyltransferase (PPT) in ubiquinone biosynthesis and boraginaceous PGTs in shikonin biosynthesis (Yazaki et al. 2002; Wang et al. 2023b).

DMAPP, geranyl diphosphate (GPP) and farnesyl pyrophosphate (FPP) were tested with DHNA (**1**) as the acceptor to explore the donor selectivity of RcDT1 (Fig. 3D). Consequently, only DMAPP could be recognized, which validated the strict prenyl donor specificity of RcDT1.

### Chemical inhibitor studies of the recombinant RcDT1

In order to obtain further information on the substrate recognition mechanism and confirm the role of *RcDT1* in the prenylation activity of *R. cordifolia* microsomes, various phenolic compounds were used for competitive analysis (Suttiyut et al. 2023). The candidate inhibitors included a MenA inhibitor Ro 48–8071 (**24**) (Dhiman et al. 2019), three phenol inhibitors for LePGT1 [homogentisic acid (**19**), catechol (**25**), and hydroquinone (**26**)] (Ohara et al. 2013), and simple naphthols (**8**, **27** – **30**) (Fig. 4A). Each of the candidate inhibitors was added to the reaction mixture at the same concentration of the substrate, and the effect on the activity of recombinant RcDT1 was examined. The competitive analyses using DHNA (**1**) or PHBA (**9**) as the substrate produced nearly identical results, hence only the results for DHNA (**1**) were shown. As seen in Fig. 4B, the MenA inhibitor Ro 48–8071 (**24**) exhibited no inhibition effect on the prenylation of DHNA (**1**) by RcDT1. Among the LePGT1 inhibitors, only catechol (**25**) showed a moderate inhibitory effect, which could reduce the activity of RcDT1 by 50.9 %. These results highlighted the difference of MenA homologs, boraginaceous PGTs, and RcDT1 in the substrate recognition mechanism. Simple naphthols 1,4-dihydroxynaphthalene (1.4-DHN, **8**) and 1,2-DHN (**27**) showed strong inhibition (5.7 % and 0 % activities of control reaction, respectively), and the prenylation of **8** or **27** was not detected. Considering that the prenylation of both DHNA (**1**) and PHBA (**9**) were strongly inhibited by simple naphthols but not phenols, it could be confirmed that RcDT1 primarily recognized naphthoic acids, and benzoic acids were prenylated through promiscuous recognition.

**Figure 4.**
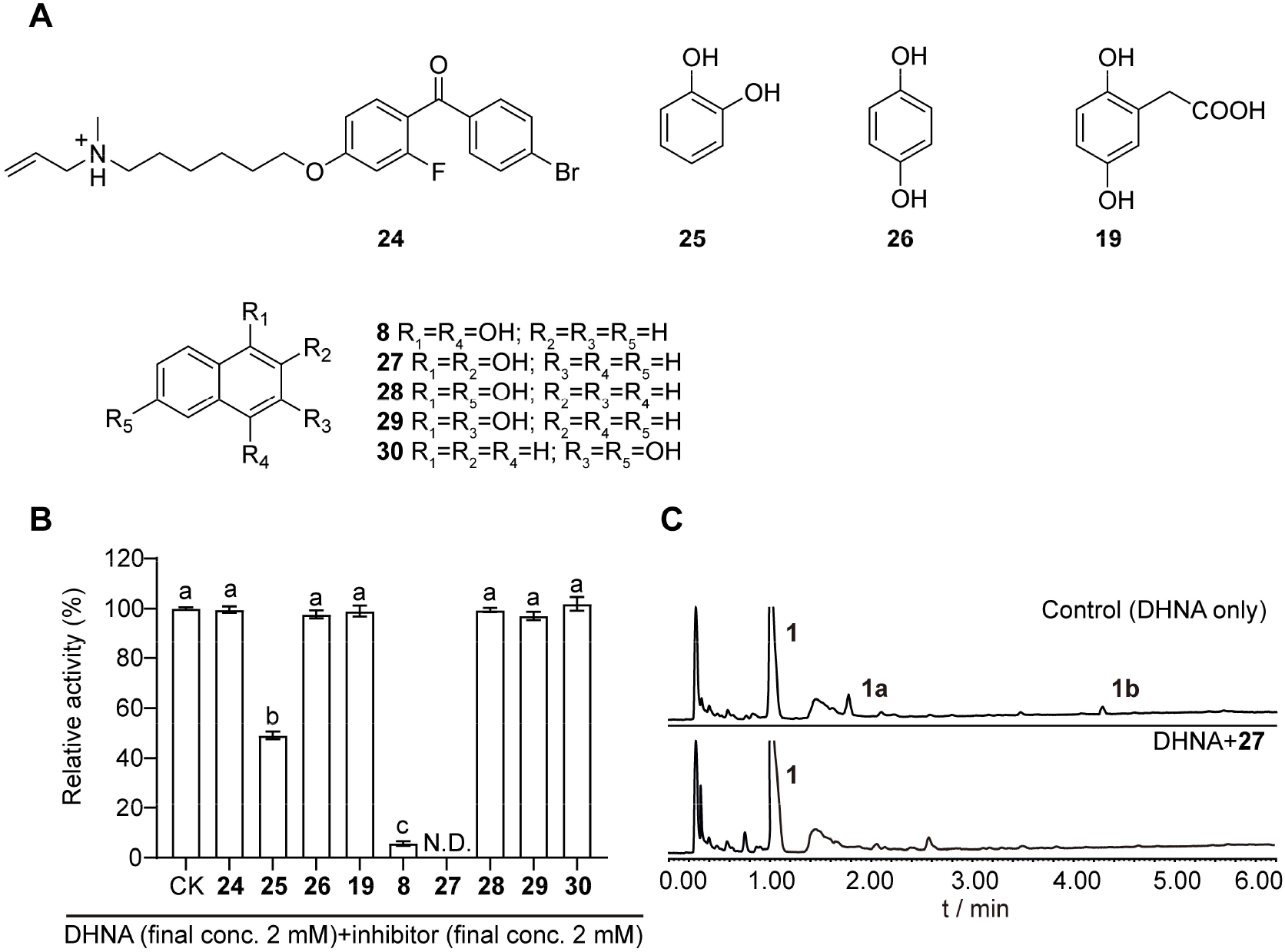
Influence of aromatic compounds on the prenylation activities of recombinant RcDT1 and *R. cordifolia* microsomes. A, Structures of tested aromatic compounds in chemical inhibitor studies. B, Relative prenylation activities for DHNA (**1**) by recombinant RcDT1 with different inhibitors. One-way ANOVA is used to compare the relative activity. Different lowercase letters indicate significant differences (*P*<0.05). N.D., not detected. C, UPLC traces for the reactions containing DHNA (**1**) with *R. cordifolia* microsomes in the presence or absence of cofactors 1,2-dihydroxynaphthalene (**27**).

The best inhibitor of RcDT1, 1,2-DHN (**27**), was then added to the reaction assay of *R. cordifolia* microsomes for competitive analysis. Like the recombinant RcDT1, 1,2-DHN (**27**) completely inhibited the prenylation catalyzed by *R. cordifolia* microsomes (Fig. 4C). This result validated that RcDT1 was responsible for the prenylation catalyzed by *R. cordifolia* microsomes.

### Biochemical properties of the recombinant RcDT1

Previous studies have shown that PT activity in plants was strongly affected by divalent cations, temperature and pH (Wang et al. 2023b). Further investigations using DHNA (**1**) as the prenyl acceptor and DMAPP as the prenyl donor revealed that the optimum reaction temperature of recombinant RcDT1 was 30 ℃, and more than 60% of the activity was maintained at a temperature up to 50 ℃ (Fig. 5A). The analysis of the enzyme activity within the pH range of 6.5 to 10.5 revealed that the optimal pH value was about 8.5, and the activity decreased rapidly at pH above 9.0 (Fig. 5B). The activity of recombinant RcDT1 was observed to decrease in the order of Mg^2+^>Fe^2+^>Mn^2+^>Co^2+^>Cu^2+^. Ca^2+^, Ni^2+^, and Zn^2+^ did not lead to the production of CPNQ (**1a**) and PNQ (**1b**). No product was detected in the reaction buffer after the addition of EDTA. (Fig. 5C).

**Figure 5.**
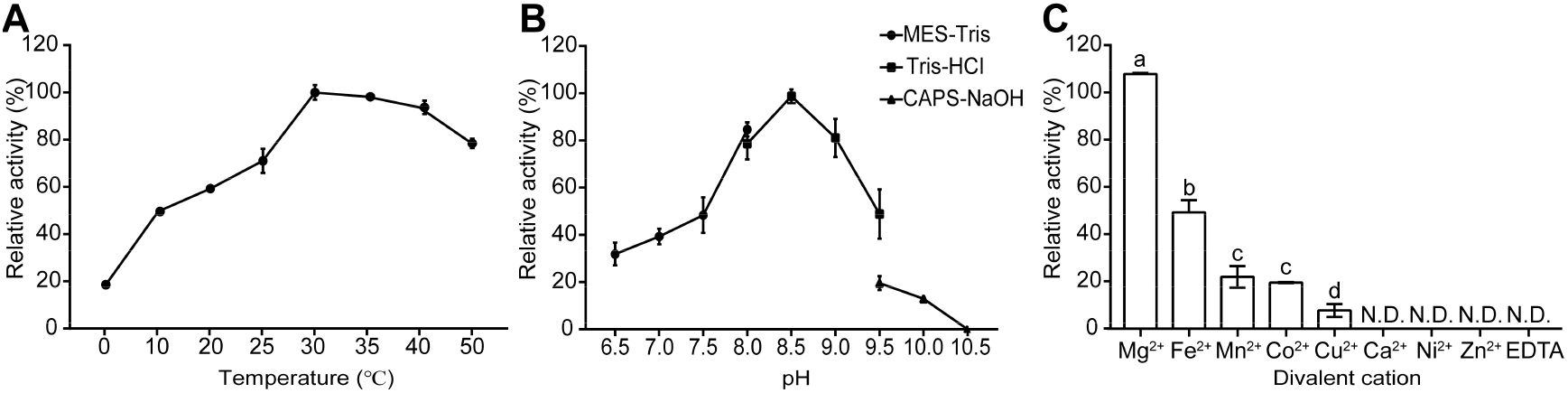
Biochemical properties of the recombinant RcDT1. A, The effects of different temperatures on the activity of recombinant RcDT1. B, PH dependences of recombinant RcDT1. C, The effects of different divalent cations on the activity of recombinant RcDT1. One-way ANOVA was used to compare the relative activity. Different lowercase letters indicate significant differences (*P*<0.05). N.D., not detected.

In order to further understand the preference of recombinant RcDT1 to prenyl acceptors, the apparent *Km* values of all substances that can be catalyzed by recombinant RcDT1 were measured with the exception of 2-hydroxy-1-naphthoic acid (**4**), 3-hydroxy-2-naphthoic acid (**5**) and orsellinic acid (**13**), which were not measured due to the low conversion rate. The results showed that DHNA (**1**) was the most suitable substrate (*Km* 26.86 ± 5.52 *μ*M). The apparent *Km* values of RcDT1 for **2**, **3**, **9**, **10**, **11** and **12** were calculated as 59.23 ± 11.20 *μ*M, 46.07 ± 6.75 *μ*M, 54.43 ± 11.41 *μ*M, 78.46 ± 19.21 *μ*M, 128.8 ± 17.31 *μ*M and 83.17 ± 11.59 *μ*M, respectively. The apparent *Km* value for DMAPP was calculated as 79.49 ± 5.93 *μ*M (Table 1).

**Table 1:**
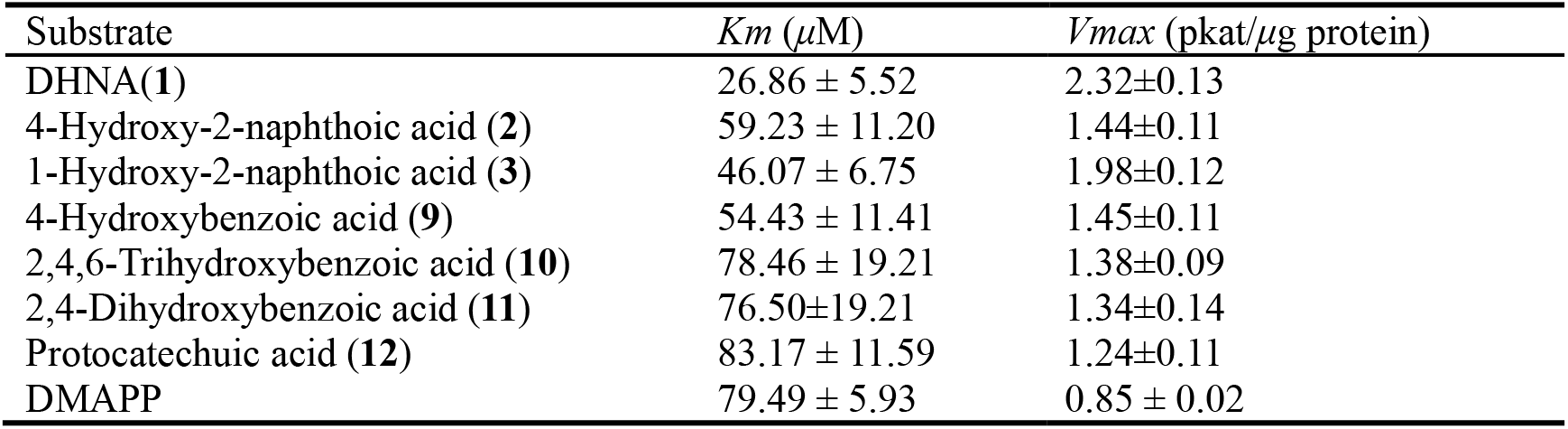
Kinetic parameters of RcDT1.

### Expression profile and subcellular localization of *RcDT1*

Analysis of the organ-specific expression of *RcDT1* in intact *R. cordifolia* plants by quantitative RT-PCR showed that the expression in roots was significantly higher than that in stems and leaves (Fig. 6A). This result is consistent with that the AQ content in the roots of *R. cordifolia* is much higher than that in other tissues (Xu et al. 2014).

**Figure 6.**
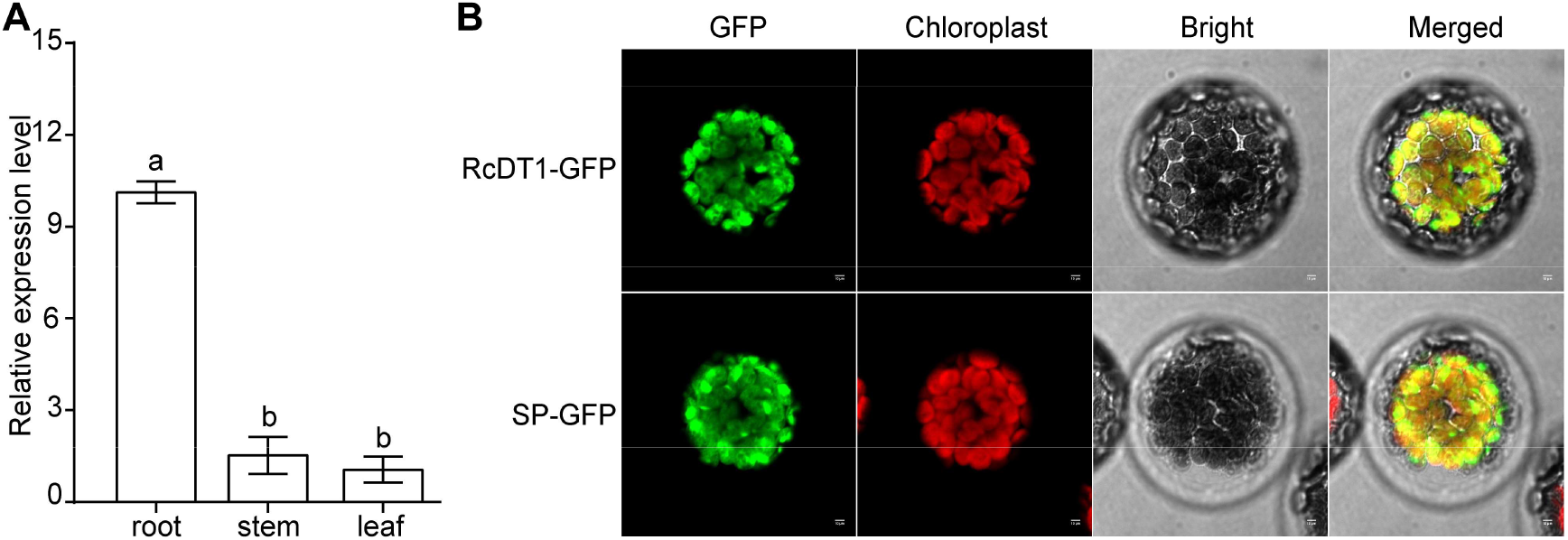
Organ-specific gene expression and subcellular localization of RcDT1. A, Expression of *RcDT1* in different organs of *R. cordifolia*. All experiments were performed in triplicate, each being repeated at least three times. The vertical bars represent the standard error of the means of independent replicates. One-way ANOVA was used to compare the expression difference between tissues. Different lowercase letters indicate significant differences (*P*<0.05). B, Subcellular localizations of full-length RcDT1 and the signal peptide (the first 72 amino acids at the N terminus) of RcDT1. RcDT1-GFP, and RcDT1SP-GFP are introduced into *A. thaliana* protoplasts. Chloroplasts are revealed by red chlorophyll autofluorescence. SP, Signal peptide. Scale bars = 10 *μ*m.

To assess the subcellular localization of RcDT1 *in planta*, full-length RcDT1 or its first 72 amino acids representing the signal peptide were fused with the green fluorescent protein (GFP) and introduced into *A. thaliana* protoplasts. Confocal microscopy of the transformed protoplasts showed that the fluorescence pattern of chloroplast was similar to that of GFP co-localized with RcDT1 or its signal peptide, indicating chloroplast localization of both chimeric proteins (Fig. 6B). The results of subcellular localization suggest that RcDT1 functions in plastids of *R. cordifolia*, consistent with the plastid localization of the MEP pathway that provides DMAPP to constitute ring C of AQs in rubiaceous plants (Han et al. 2001).

### Functional validation of RcDT1 homologs in other rubiaceous plants

Considering that AQs found in Rubiaceae are all formed through the shikimate pathway, we attempt to verify whether homologs of RcDT1 in other rubiaceous plants have similar functions as in *R. cordifolia*. The polypeptide sequence of *RcDT1* was employed as a query to retrieve the genome databases of *C. arabica* and *C. eugenioides* (two classical examples of Rubiaceae), and the transcriptome database of *M. officinalis* (an AQ-rich medicinal plant widely cultivated in the Lingnan region of southern China) independently (Wang et al. 2021b). Regarding *C. arabica* and *C. eugenioides*, the search for genes with identities greater than 60% to *RcDT1* yielded 4 and 2 homologs, respectively. After removing highly similar genes, three homologs termed *CaDT1*, *CaDT2*, and *CeDT1* were selected for following studies (Table S1). As for *M. officinalis*, *MoDT1*, the only gene with significant homology to *RcDT1* (73.79%), was investigated in follow-up experiments.

Truncated *CaDT1*, *CaDT2*, *CeDT1*, and *MoDT1* genes with a 72 amino acid deletion from the start codon were synthesized and expressed in *P. pastoris*. The investigation for the prenylation activity was carried out using DHNA (**1**) or PHBA (**9**) as the substrate with recombinant enzymes in the presence of DMAPP and Mg^2+^. As a result, recombinant CaDT1 and MoDT1 could prenylate DHNA (**1**) and PHBA (**9**) in the same manner as recombinant RcDT1 (Fig. S2). This suggests that the prenylation step of AQ biosynthesis in Rubiaceae is catalyzed by RcDT1 and its homologs. Recombinant CaDT2 did not exhibit any prenylation activity toward DHNA (**1**) and PHBA (**9**). Interestingly, recombinant CeDT1 could only accept PHBA (**9**) but not DHNA (**1**) as the prenyl acceptor (Fig. S3). The promiscuous recognition of DHNA (**1**) and PHBA (**9**) by RcDT1 and its homologs support the view that DHNA PTs and PHBA PTs share a common evolutionary origin in Rubiaceae.

### Phylogenetic analysis of *RcDT1*

*RcDT1* was used to query the NCBI Gene database (https://www.ncbi.nlm.nih.gov/gene) and OneKP database (https://db.cngb.org/onekp/search) to obtain homologous sequences from Rubiaceae. A maximum-likelihood tree was constructed to analyze the evolutionary relationships of *RcDT1* and its homologs with the other plant aromatic PTs (Fig. 7). *RcDT1* and its homologs turned out to form a new branch having a common origin with the PGTs for shikonin formation in Boraginaceae and PPTs for ubiquinone formation. Correspondingly, *RcDT1* and its homologs in Rubiaceae could recognize PHBA (**1**) as the substrate. Therefore, it can be concluded that DHNA/PHBA DTs in Rubiaceae have evolved via recruitment from the ubiquinone biosynthetic pathway. On the other hand, the rubiaceous branch in the tree was clearly divergent from that of the MenA homologs for phylloquinone biosynthesis. This was verified by the result that recombinant RcDT1 and MenA were different in substrate recognition and prenylation mechanisms. These phylogenetic analyses suggested that DHNA-prenylation activities evolved convergently in distant clades of the UbiA superfamily.

**Figure 7.**
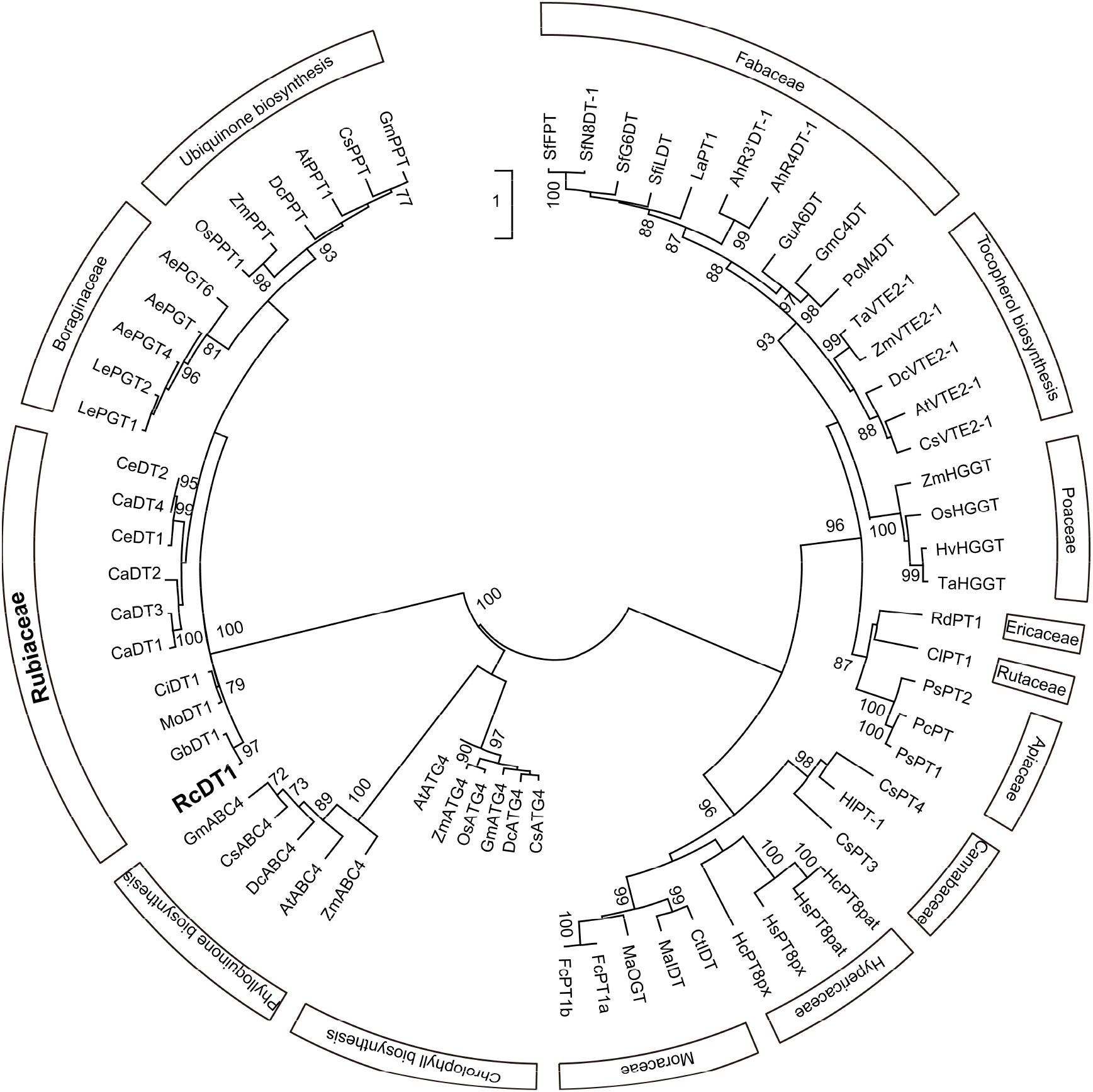
Phylogenetic analysis of *RcDT1*. The phylogenetic relationship between the *RcDT1* protein and the related prenyltransferases. The tree was constructed using the maximum-likelihood method applying 1000 bootstrap replicates. Nodes with bootstrap values >70 are indicated by numbers on the branch. The scale bar represents 1 amino acid substitutions per site. SfFPT (*Sophora flavescens*; KC513505); SfN8DT-1 (*S. flavescens*; AB325579); LaPT1 (*Lupinus albus*; JN228254); SfG6DT (*S. flavescens*; AB604224); SfiLDT (*S. flavescens*; AB604223); GuA6DT (*Glycyrrhiza uralensis*; KJ123716); GmC4DT (*Glycine max*; LC140927); PcM4DT (*Psoralea corylifolia*; MH626730); AhR3’DT-1 (*Arachis hypogaea*; KY565245); AhR4DT-1 (*A. hypogaea*; KY565244) TaVTE2-1 (*Triticum aestivum*; DQ231056); ZmVTE2-1 (*Zea mays*; EU973221); AtVTE2-1 (*A. thaliana*; DQ231056); CsVTE2-1 (*Citrus sinensis*; XM_006474143); DcVTE2-1 (*Daucus carota*; XM_017398463); HvHGGT (*Hordeum vulgare*; AY222860); TaHGGT (*Triticum aestivum*; AY222861); OsHGGT (*Oryza sativa*; AY222862); ZmHGGT (*Z. mays*; XM_008661550); ClPT1 (*Citrus limon*; AB813876); RdPT1 (*Rhododendron dauricum*; LC381857); PsPT2 (*Pastinaca sativa*; KM017084); PcPT (*Petroselinum crispum*; AB825956); PsPT1 (*P. sativa*; KM017083); CsPT3 (*Cannabis sativa*; BK010680); CsPT4 (*C. sativa*; BK010648); HlPT-1 (*Humulus lupulus*; AB543053); HcPT8pat (*Hypericum calycinum*; MH461103); HcPT8px (*H. calycinum*; MH461102); HsPT8px (*Hypericum sampsonii*; MH461100); HsPT8pat (*H. sampsonii*; MH461101); CtIDT (*Cudrania tricuspidata*; KM262660); MaIDT (*Morus alba*; KM262659); FcPT1a (*Ficus carica*; LC369744); FcPT1b (*F. carica*; LC369745); OsATG4 (*O. sativa*; EF432576); ZmATG4 (*Z. mays*; NM_001148732); AtATG4 (*A. thaliana*; NM_115041); DcATG4 (*D. carota*; XM_017388300); CsATG4 (*C. sinensis*; XM_006481001); GmATG4 (*G. max*; NM_001252704); AtABC4 (*A. thaliana*; NM_001124046.2); CsABC4 (*C. sinensis*; XM_025093178); GmABC4 (*G. max*; XM_003532557); ZmABC4 (*Z. mays*; NM_001158698); DcABC4 (*D. carota*; AY222861); RcDT1 (*R. cordifolia*; OR493119); CaDT1 (*C. arabica*; XM_027266276); CaDT2 (*C. arabica*; XM_027266749); CaDT3 (*C. arabica*; XM_027269080); CaDT4 (*C. arabica*; XM_027124411); CeDT1 (*C. eugenioides*; XM_027313870); CeDT2 (*C. eugenioides*; XM_027313873); CiDT1 (*Carapichea ipecacuanha*; BQEQ_2057440); GbDT1 (*Galium boreale*; WQRD_2015494); AePGT (*A. euchroma*; DQ397513); AePGT4 (*A. euchroma*; KT991522); LePGT2 (*Lithospermum erythrorhizon*; AB055079); LePGT1 (*L. erythrorhizon*; AB055078); AePGT6 (*A. euchroma*; KT991524); DcPPT (*D. carota*; XM_017367441); AtPPT1 (*A. thaliana*; NM_118497); GmPPT (*G. max*; XM_006602661); CsPPT (*C. sinensis*; XM_015532016); OsPPT1 (*O. sativa*; AY222861); ZmPPT (*Z. mays*; NM_001155086). The sequence MoDT1 from *M. officinalis* can be found in Table S3. Table 1 Kinetic parameters of RcDT1.

## Discussion

Alizarin-type AQs are mainly distributed in lamiids clade (Bignoniaceae, Gesneriaceae, Lamiaceae, Rubiaceae, Scrophulariaceae, and Verbenaceae), as well as in Amaranthaceae, Euphorbiaceae, and Juglandaceae (Thomson 1991; Wang et al. 2023a). Among these families, Rubiaceae is a rich source of alizarin-type AQs (Martins and Nunez 2015). And rubiaceous plants accumulating alizarin-type AQs are all medicinal plants (Wang et al. 2023a). Rubiaceous plants biosynthesize AQs through the combination of shikimate pathway providing the DHNA moiety and MEP pathway providing the prenyl (isoprenoid) chain (Han et al. 2001). The same route has also been used to synthesize alizarin-type AQs in Bignoniaceae and Verbenaceae (Fig. 1) (Burnett and Thomson 1968a and 1968b). The key step in the coupling of the two pathways is catalyzed by a membrane-bound aromatic PT belonging to the UbiA superfamily. However, its genetic information, enzymatic characteristics, reaction mechanism, resulting intermediate(s), and evolution remain obscure. In this study, we started by establishing cell suspension cultures of *R. cordifolia* in which a DHNA-DT activity was detected. Then a candidate PT gene belonging to UbiA superfamily, *RcDT1*, was discovered to account for the prenylation activity. The prenylation of DHNA was accompanied by spontaneous oxidation and decarboxylation, resulting in the formation of CPNQ (**1a**) and deoxylapachol (**1b**). The recombinant RcDT1 primarily recognized naphthoic acids with hydroxyl(s) in the carboxyl-bearing ring, and the benzoic acids with 4-hydroxyl group were also prenylated through promiscuous recognition. Chemical inhibitor studies confirmed the role of *RcDT1* in the prenylation catalyzed by *R. cordifolia* suspension cell. The plastid localization and root-specific expression of *RcDT1* further proved its participation in the biosynthesis of AQs. When the study was expanded to other rubiaceous plants (*C. arabica*, *C. eugenioides*, and *M. officinalis*), the homologs of *RcDT1* also exhibited similar prenylation activities toward DHNA and PHBA, which validated the role of *RcDT1* and its homologs in the biosynthesis of AQs from Rubiaceae. Finally, the phylogenetic analyses confirmed that DHNA/PHBA DTs in Rubiaceae have evolved via recruitment from the ubiquinone biosynthetic pathway.

Compared with the identified DHNA PTs homologous to MenA, RcDT1 differed significantly in reaction mechanism. The tracer studies proved that the prenylation of DHNA by MenA occurred at C-2 position and required replacement of the carboxyl with the isoprenoid side chain (Baldwin et al. 1973). A carbocation mechanism was postulated in which prenylation and decarboxylation occurred in a single active site (Meganathan and Kwon 2009). This is supported by the fact that no intermediate or two separate reaction steps are observed for MenA catalyzed prenylation. In contrast, the prenylation of DHNA by RcDT1 occurred at C-3 position and it was uncoupled from the decarboxylation. As in MenA-catalyzed prenylation, the introduction of dimethylallyl moiety by RcDT1 was accompanied by a spontaneous naphthol to naphthoquinone oxidation to produce CPNQ (**1a**). The formation of β-keto structure may facilitate the decarboxylation to generate PNQ (**1b**). A similar decarboxylation step was proposed in the laccase-catalyzed synthesis of 1,4-naphthoquinone-2,3-bis-sulfides from DHNA (Wellington et al. 2012). The main biological significance of this stepwise mechanism is that it leads to the branch of the biosynthetic pathway towards AQs and NHQs in Rubiaceae (Fig. 1). Specifically, if **1a** was further methyl-esterified to form 2-methoxycarbonyl-3-prenyl-1,4-naphthoquinone, its decarboxylation and subsequent cyclization to form AQs were prevented and the metabolic flux was shifted to NHQs (Fig. 1). When incubated in the reaction buffer without proteins, 2-methoxycarbonyl-3-prenyl-1,4-naphthoquinone was observed to spontaneously cyclize to form mollugin, a signature naphthoquinone of *R. cordifolia* (Fig. S4). In contrast, the decarboxylation of **1a** to form **1b** triggered the subsequent cyclization and enabled entry into alizarin-type AQs (Fig. 1). In addition to Rubiaceae, alizarin-type AQs are also found in Bignoniaceae and Verbenaceae, where they are accompanied by the structure-related characteristic constituent lapachol (Hussain et al. 2007). Therefore, it can be inferred that AQs and lapachol are both biosynthesized through the shikimate pathway, and PNQ (i.e. deoxylapachol, **1b**) is also the immediate precursor of lapachol and related quinones (Fig. 1).

The above differences in reaction mechanism, together with the phylogenetic clustering, strongly suggest that the DHNA-prenylation activities evolved convergently from different ancestral UbiA progenitors in ubiquinone and phylloquinone biosynthesis, respectively. This evolutionary trajectory is in line with previous reports describing the convergent evolution of coumarin C-PTs in Apiaceae and Moraceae (Munakata et al. 2020), coumarin O-PTs in Apiaceae and Rutaceae (Munakata et al. 2021), as well as flavonoid and stilbene PTs in Moraceae and Fabaceae (Wang et al. 2014; Yang et al. 2018; Zhong et al. 2018). These recurring neofunctionalization events could be due to a number of factors (Lou et al. 2022). Firstly, UbiA superfamily is a large gene family and its members are widely involved in the biosynthesis of both primary and specialized metabolites in microbes, plants, and animals. Diverse substrate specificities and expression profiles of UbiA PTs increase opportunities of neofunctionalization. Secondly, substrate promiscuity was repeatedly found in UbiA PTs of specialized metabolism (Chen et al. 2013), which served as the starting point for neofunctionalization (Weng et al. 2012). Finally, the end product of the pathway containing UbiA PTs may have abundant ecological functions in multiple niches, which forces the mutations to be preserved.

As the first pathway-specific enzyme of alizarin-type AQ biosynthesis, *RcDT1* independently evolved from the highly conserved PPT of ubiquinone biosynthesis. Both its promiscuous recognition of the benzoic acids with 4-hydroxyl group and phylogenetic clustering support this conclusion. Interestingly, *CeDT1*, a homolog of *RcDT1* in *C. eugenioides*, lost the DHNA-prenylation activity but retained PHB-prenylation activity, confirming the evolutionary relevance of the two functions in Rubiaceae. A similar evolutionary event took place in Boraginaceae, wherein PGTs of shikonin biosynthesis evolved independently from PPTs. Boraginaceae and Rubiaceae are classified as Boraginales and Gentianales, respectively, both orders are core groups in the lamiids clade. The taxonomic neighborhood of the two orders explained the relative high amino acid identities of RcDT1 with boraginaceous PGTs. Both RcDT1 and boraginaceous PGTs function as the first pathway-specific enzyme of quinone biosynthesis. And similar cyclization mechanisms were observed in the subsequent biosynthetic reactions of these two classes of quinones (Fig. 1). In cell cultures of *R. cordifolia* and *L. erythrorhizon*, both AQs and NQs formation were repressed by irradiation of blue light. Taken together, it could be drawn that neofunctionalization events of PPTs in Rubiaceae and Boraginaceae may lead to taxon-specific metabolic pathways to form AQs and NQs. And homology might be predicted for subsequent steps in AQ and NQ biosynthesis. However, the remaining reaction steps in shikimate pathway, especially the cyclization, still remain to be fully elucidated.

Excluding AQs, shikonin and lapachone derivatives, shikimate pathway has been found to form NQs such as chimaphilin in the distantly-related plants from Pyroleae (Fig. 1) (Meyer et al. 2021). A deeper understanding is needed in terms of achieving analogous chemicals by convergent evolution of this pathway in different plants.

In conclusion, RcDT1, the first pathway-specific PT in the biosynthesis of alizarin-type AQs, was identified from *R. cordifolia*. In contrast to the known DHNA PTs, the prenylation of DHNA by RcDT1 proceeds in a stepwise manner, yielding two products via direct prenylation and decarboxylation. Phylogenetic analysis further proved that RcDT1 and its rubiaceous homologs evolved convergently from ancestral UbiA progenitors in ubiquinone biosynthesis. As the branch point enzyme in shikimate pathway, RcDT1 and its homologs opened the biosynthesis of alizarin-type AQs and NQs not only in Rubiaceae, but possibly in other lamiids plants as well. Thus the evolution of RcDT1 provides useful guidance for identifying additional and evolutionarily varied PTs which enable entry into quinones derived from shikimate pathway. Moreover, the identification and characterization of RcDT1 will have profound implications for understanding the biosynthetic process of the AQ/NQ ring derived from shikimate pathway.

## Materials and methods

### General experimental procedures

^1^H and ^13^C NMR spectra were recorded on Bruker DRX 500 spectrometer. The observed chemical shift values were reported in ppm. The UPLC and LC-MS analyses of the enzymatic products were performed as previously described with the exception that the solvent system comprised acetonitrile (containing 0.1% formic acid, A) and water (containing 0.1% formic acid, B) at 0.5 mL min^-1^ with the following gradient program: 0 min, 30% A; 6 min, 60% A; 6.5 min, 100% A; 8.5 min, 100% A (Wang et al., 2019). A Waters Acquity UPLC-I Class system (Waters Co., Milford, MA, USA) was equipped with a BEH Phenyl column (2.1 × 50 mm, 1.7 *μ*m), and the absorbance at 254 nm was measured. For the isolation of the enzymatic product, the same approach as previously described was employed (Wang et al. 2019).

### Plant materials and chemicals

Young leaves of *R. cordifolia* were collected on the campus of China Academy of Chinese Medical Sciences in August, 2018, and used as explants to initiate calli. The initiation and maintaince of the calli were performed as previously described with the exception that MS medium supplemented with 1.0 mg·L^-1^ naphthalene acetic acid and 0.5 mg·L^-1^ 6-benzylaminopurine was used (Yin et al. 2013).

Dimethylallyl diphosphate (DMAPP) was synthesized according to a previously described method (Davisson et al. 1985), other chemical reagents were purchased from Sigma-Aldrich (St. Louis, MO, USA) and Macklin (Shanghai, CHN).

### RNA sequencing and bioinformatic processing

The cDNA libraries of *R. cordifolia* cultured cells were sequenced on an Illumina HiSeq 2000 platform, and the raw sequencing data were deposited in the National Center for Biotechnology Information’s Short Read Archive (SRA) under accession number PRJNA1016061. The sequencing data were analyzed following the process described in a previous study (Wang et al. 2019). The raw sequencing data of *M. officinalis* (SRA accession no. SRP258544) were processed as above and used for homology search.

### Total RNA extraction and RT-qPCR analysis

Total RNA from different tissues and cultured *R. cordifolia* cells were extracted using E.Z.N.A.^TM^ Plant RNA kit (Omega Bio-tek Inc., Doraville, GA, USA) following the manufacturer’s instructions. First-strand cDNAs were synthesized with the PrimerScript First Strand cDNA Synthesis Kit at the same time [Takara Biomedical Technology (Beijing) Co., Ltd., Beijing, CHN]. RT-qPCR was performed using the TB Green^TM^ *Premix Ex Taq*™ II [Takara Biomedical Technology (Beijing) Co., Ltd., Beijing, CHN] and an Applied Biosystems 7500 realtime instrument. The primers used are listed in Table S2. The *RcActin* was used as the endogenous control to normalize expression data. These PCRs were conducted using an amplification programme consisting of initial denaturation at 95°C for 2 min followed by 40 cycles of denaturation at 95°C for 10 s, and elongation at 60°C for 30 s. Amplification of the target sequences was confirmed by sequencing. The expression level of each gene was quantified using 2^-ΔΔCt^. At least three independent experiments were performed for each analysis.

### cDNA cloning and heterologous expression in yeast

AtABC4 (GenBank accession no. NP_001117518.1), LePGT1 (GenBank accession no. BAB84122.1), AePGT6 (GenBank accession no. ANC67959.1), and SfN8DT-1 (GenBank accession no. BAG12671.1) were used as queries to retrieve the transcriptome database of *R. cordifolia* cell suspension cultures. As a result, six candidate genes numbered as *RcDT1 – 6* were selected for further analysis. Specific primers were designed to obtain the complete coding sequence of *RcDT1 – 6* from the cDNA of cultured *R. cordifolia* cells (Table S1). Using the complete coding sequences as the templates, primers were designed to amplify truncated *RcDT1* (with an 24/48/72-amino acid deletion from the start codon) and truncated *RcDT2 – 6* (with an 24-amino acid deletion from the start codon). Complete and truncated PT sequences were subcloned into the *EcoR* I/*Kpn* I restriction sites of the pPink-HC plasmid, and the plasmid-construction were verified by sequencing.

The expression vectors were introduced into the yeast PichiaPink™ Strain 2 using Frozen-EZ Yeast Transformation II™ (Zymo Research Co., Irvine, CA, USA). Recombinant strains were initially cultured in BMGY liquid medium at 30 °C for about 24 h. The cells were then harvested and resuspended into BMMY liquid medium to induce recombinant proteins expression. The induction lasted 5 days at 30 °C, and methanol was added to the final concentration of 0.5% on time every day.

### *In vitro* enzyme activity assay

The prenylation activity was assayed for recombinant yeasts and cultured *R. cordifolia* cells respectively. The collected yeast cells were broken with a APV2000 continuous high-pressure crusher (AxFlow; Stockholm; SWE). The cultured cells of *R. cordifolia* were ground in a mortar. The microsomal proteins were prepared by differential centrifugation as described previously (Wang et al. 2014). All these procedures were performed at 4 ℃. The total protein concentration was determined by the Bradford method.

The reaction mixture (200 *μ*L) for determining enzyme activity contained 50 mM Tris-HCl (pH 8.5), 5 mM MgCl_2_, 200 *μ*M the prenyl acceptors, 400 *μ*M DMAPP, and 0.5 mg microsome protein. The reaction mixtures were incubated at 30 °C for 30 min, and the reaction was terminated by the addition of 600 *μ*L of acetonitrile. The protein was removed by centrifugation at 15000 × *g* for 20 min, and the reaction was analyzed by UPLC-UV/ESI MS.

### Biochemical properties of the recombinant enzymes

With DHNA as the prenyl acceptor and DMAPP as the prenyl donor, the biochemical characteristics of the recombinant enzyme were analyzed by calculating the yield of the main product. To investigate the optimal pH, the enzyme reactions were performed in reaction buffers with pH values in the range of MES-Tris (pH 6.5-8.0), Tris-HCl (pH 8.0-9.5), and CAPS-NaOH (pH 9.5-10.5) at 30 °C for 30 min. To investigate the optimal temperature, the reaction mixtures were incubated at eight different temperatures (0 °C to 50 °C) in 50 mM Tris-HCl buffer (pH 8.5). To test the requirement of *RcDT1* activity for divalent cations, the reaction mixtures were incubated with 5 mM MgCl_2_, CaCl_2_, FeCl_2_, NiCl_2_, MnCl_2_, CuCl_2_, ZnCl_2_, CoCl_2_ and EDTA respectively in 50 mM Tris-HCl buffer (pH 8.5) at 30 °C for 30 min. Relative activities of recombinant RcDT1 with prenyl diphosphate of different chain lengths were analyzed with DMAPP (C5), GPP (C10), and FPP (C15) respectively at 30 °C for 30 min.

The apparent *Km* values for different prenyl acceptors were determined by incubating 200 *μ*g of recombinant yeast microsomes with various concentrations of prenyl acceptors (12-400 *μ*M) and a fixed concentration of DMAPP (400 *μ*M) in 200 *μ*L reaction mixtures at 30 °C. The apparent *Km* values for DMAPP were determined using various concentrations of DMAPP (12-400 *μ*M) and a fixed concentration of DHNA (400 *μ*M) under the same conditions. For each prenyl acceptors or DMAPP, the incubation time was controlled respectively so that the reaction was under 10% complete. Kinetic constants were calculated based on Michaelis-Menten kinetics using GraphPad Prism 8 (GraphPad Software Inc., San Diego, CA, USA).

### Structure identification of enzymatic products

The assays for the isolation of the enzymatic products (15-20 ml) contained 50 mM Tris-HCl (pH 8.5), 5 mM MgCl_2_, 2 mM the prenyl acceptors, 4 mM DMAPP, and 30 mg of the recombinant RcDT1 microsome. The reaction mixtures were incubated at 30 °C for 12 h and subsequently extracted three times with ethyl acetate. After evaporation of the solvent under vacuum, the residues were dissolved in methanol and purified by reverse-phase semi-preparative HPLC. The products were analyzed by MS, and ^1^H and ^13^C NMR spectroscopy, which yielded the following results.

3-prenyl-4-hydroxy-2-naphthoic acid (**2a**): TOF-MS, *m*/*z*: 255.1 [M-H]^-^; ^1^H NMR (500 MHz, acetone-*d*_6_): *δ* 8.28 (1H, *d*, *J* = 8.5 Hz, H-5), 8.05 (1H, br. *s*, H-1), 7.93 (1H, *d*, *J* = 8.5 Hz, H-8), 7.56–7.59 (1H, *m*, H-7), 7.50–7.53 (1H, *m*, H-6), 5.25 (1H, br. *t*, *J* = 7.0 Hz, H-2’), 3.96 (2H, *d*, *J* = 7.0 Hz, H-1’), 1.78 (3H, br. *s*, H-4’), 1.64 (3H, br. *s*, H-5’) (Fig. S4); ^13^C NMR (125 MHz, acetone-*d*_6_): *δ* 169.78 (COOH), 150.99 (C-4), 132.81 (C-9), 131.94 (C-3’), 129.52 (C-7), 127.84 (C-8), 127.64 (C-10), 126.95 (C-6), 126.64 (C-2), 124.62 (C-5), 123.98 (C-2’), 122.82 (C-3), 122.36 (C-1), 26.14 (C-1’), 25.90 (C-5’), 18.15 (C-4’) (Fig. S5). The chemical structure of **2a** was determined by comparison of the spectroscopic data with the substrate **2** (Liang et al. 2014).

4-prenyl-1-hydroxy-2-naphthoic acid (**3a**): TOF-MS, *m*/*z*: 255.1 [M-H]^-^; ^1^H NMR (500 MHz, acetone-*d*_6_): *δ* 8.38 (1H, *d*, *J* = 8.5 Hz, H-8), 7.97 (1H, *d*, *J* = 8.0 Hz, H-5), 7.75 (1H, br. *s*, H-3), 7.62–7.64 (1H, *m*, H-6), 7.50–7.53 (1H, *m*, H-7), 5.37 (1H, br. *t*, *J* = 7.0 Hz, H-2’), 3.68 (2H, *d*, *J* = 7.0 Hz, H-1’), 1.81 (3H, br. *s*, H-4’), 1.73 (3H, br. *s*, H-5’) (Fig. S6); ^13^C NMR (125 MHz, acetone-*d*_6_): *δ* 174.60, 160.14, 136.32, 132.63, 130.52, 129.18, 127.49, 126.30, 125.59, 124.84, 124.79, 124.42, 108.91, 25.88, 25.66, 17.99 (Fig. S7). The chemical structure of **3a** was determined by comparison of the spectroscopic data with the substrate **3** (Wang and Gevorgyan 2015).

3-prenyl-4-hydroxybenzoic acid (**9a**): TOF-MS, *m*/*z*: 205.1 [M-H]^-^; ^1^H NMR (500 MHz, acetone-*d*_6_): *δ* 7.81 (1H, *d*, *J* = 2.0 Hz, H-2), 7.75 (1H, *dd*, *J* = 8.0 Hz, 2.0 Hz, H-1), 6.91 (1H, *d*, *J* = 8.0 Hz, H-5), 5.35 (1H, br. *t*, *J* = 7.5 Hz, H-2’), 3.57 (2H, *d*, *J* = 7.5 Hz, H-1’), 1.73 (6H, br. *s*, H-4’, 5’) (Fig. S8); ^13^C NMR (125 MHz, acetone-*d*_6_): *δ* 167.64 (COOH), 160.11 (C-4), 133.10 (C-3’), 132.34 (C-1), 130.14 (C-6), 128.84 (C-3), 123.06 (C-2’), 122.66 (C-1), 115.39 (C-5), 28.79 (C-1’), 25.90 (C-5’), 17.83 (C-4’) (Fig. S9) (Abraham and Arfmann 1990).

5-prenyl-4-hydroxysalicylic acid (**11a**): TOF-MS, *m*/*z*: 221.1 [M-H]^-^; ^1^H NMR (500 MHz, acetone-*d*_6_): *δ* 7.61 (1H, *s*, H-6), 6.39 (1H, *s*, H-3), 5.32 (1H, br. *t*, *J* = 7.5 Hz, H-2’), 3.24 (2H, *d*, *J* = 7.5 Hz, H-1’), 1.72 (3H, br. *s*, H-4’), 1.70 (3H, br. *s*, H-5’) (Fig. S10); ^13^C NMR (125 MHz, acetone-*d*_6_): *δ* 173.11 (COOH), 163.20 (C-4), 162.08 (C-2),132.50 (C-3’), 132.00 (C-6), 123.68 (C-2’), 120.60 (C-5), 106.16 (C-1), 102.79 (C-3), 28.20 (C-1’), 25.91 (C-5’), 17.80 (C-4’) (Fig. S11) (Abraham and Arfmann 1990).

### Subcellular localization of *RcDT1*

The nucleotide sequence for full-length *RcDT1* or its N-terminal sequence (216 bp) were subcloned into the *BsaI*/*Eco31I* restriction sites of the pBWA(V)HS-ccdb-GLosgfp plasmid to give pBWA(V)HS-RcDT1, pBWA(V)HS-RcDT1-TP. *Arabidopsis thaliana* protoplasts were isolated and transformed as described previously with modifications (Yoo et al. 2007). The transformed protoplasts were examined with a light microscope as well as a laser confocal microscope (C2-ER; Nikon). For green fluorescent protein (GFP) detection, excitation at 488 nm and detection at 510 nm were used. For chloroplast detection, excitation at 640 nm and detection at 675 nm were used.

### In silico sequence analysis

The transmembrane regions and the transit peptides of RcDTs were predicted by DeepTMHMM (https://dtu.biolib.com/DeepTMHMM) and SignalP - 6.0 (https://services.healthtech.dtu.dk/services/SignalP-6.0/), respectively. The protein sequences were aligned using Clustal W, and phylogenetic tree was drawn by MEGA X. Phylogenetic relationships were reconstructed by the maximum likelihood method based on the JTT/+G model (five categories) and a bootstrap of 1,000 replicates. Bootstrap values were indicated in percentages (only those >70 % were presented) on the nodes. The scale bar corresponded to 1.0 estimated amino acid changes per site.

### Accession numbers

The nucleotide sequences of *RcDT1 – 6* have been deposited in the GenBank^TM^ database under the accession numbers OR493119 – OR493124, respectively.

## Acknowledgments

We would like to thank Dr. Haiyu Zhao for providing technical support.

## Author Contributions

R. Wang, L. Guo, and L. Huang conceived and designed research. R. Wang, C. Liu, and S. Wang conducted the most experiments. T. Chen performed the bioinformatics analyses. C. Lyu, C. Kang, X. Wan, and J. Guo analyzed the data. J. Guo helped discussing and designing experiments. C. Liu and R. Wang wrote the manuscript. J. Guo, and L. Guo revised the manuscript. All authors read and approved the manuscript.

## Abbreviations

AQ: anthraquinone
NQ: naphthoquinone
PHBA: 4-hydroxybenzoic acid
PGT: PHBA geranyltransferase
PPT: PHBA polyprenyltransferase
DHNA: 1,4-dihydroxy-2-naphthoic acid
CPNQ: 2-carboxyl-3-prenyl-1,4-naphthoquinone
PNQ: 3-prenyl-1,4-naphthoquinone
DMAPP: 3,3-dimethylallyl pyrophosphate
GPP: geranyl pyrophosphate
FPP: farnesyl pyrophosphate
DHN: dihydroxynaphthalene

