## Supplemental Figures 1-11 and Supplemental Tables 1-3 for "The Discovery and Characterization of 1,4-Dihydroxy-2-naphthoic Acid Prenyltransferase Involved in the Biosynthesis of Anthraquinones in *Rubia cordifolia*"

### Supplemental Data

**Figure S1.** Alignment of the encoded polypeptides of the candidate genes……………………..…..2
**Figure S2.** Comparison of the prenylation activities of recombinant RcDT1 and its rubiaceous homologs………………………………………………………………………...…………...…..3
**Figure S3.** Spontaneous cyclization of 2-methoxycarbonyl-3-prenyl-1,4-naphthoquinone...…...….3
**Figure S4.** [1]H NMR spectrum of 3-prenyl-4-hydroxy-2-naphthoic acid (**2a**) in acetone-$d_6$ (500 MHz)…………………………………………………………………………………………..5
**Figure S5.** [13]C NMR spectrum of 3-prenyl-4-hydroxy-2-naphthoic acid (**2a**) in acetone-$d_6$ (500 MHz)…………………………………………………………………………………………..5
**Figure S6.** [1]H NMR spectrum of 4-prenyl-1-hydroxy-2-naphthoic acid (**3a**) in acetone-$d_6$ (500 MHz)………………………………………………………………………………..………6
**Figure S7.** [13]C NMR spectrum of 4-prenyl-1-hydroxy-2-naphthoic acid (**3a**) in acetone-$d_6$ (500 MHz)………………………………………………………………………………..………6
**Figure S8.** [1]H NMR spectrum of 3-prenyl-4-hydroxybenzoic acid (**9a**) in acetone-$d_6$ (500 MHz)………………………………………………………………………………..………7
**Figure S9.** [13]C NMR spectrum of 3-prenyl-4-hydroxybenzoic acid (**9a**) in acetone-$d_6$ (500 MHz)………………………………………………………………………………..………7
**Figure S10.** [1]H NMR spectrum of 5-prenyl-4-hydroxysalicylic acid (**11a**) in acetone-$d_6$ (500 MHz)………………………………………………………………………………..………8
**Figure S11.** [13]C NMR spectrum of 5-prenyl-4-hydroxysalicylic acid (**11a**) in acetone-$d_6$ (500 MHz)………………………………………………………………………………..………8
**Table S1.** The amino acid residues identities between RcDT1 and its rubiaceous homologs………..9
**Table S2.** Primers used in this study……………………………………………………………..10
**Table S3.** The sequences used in this study……………………………………………………...11

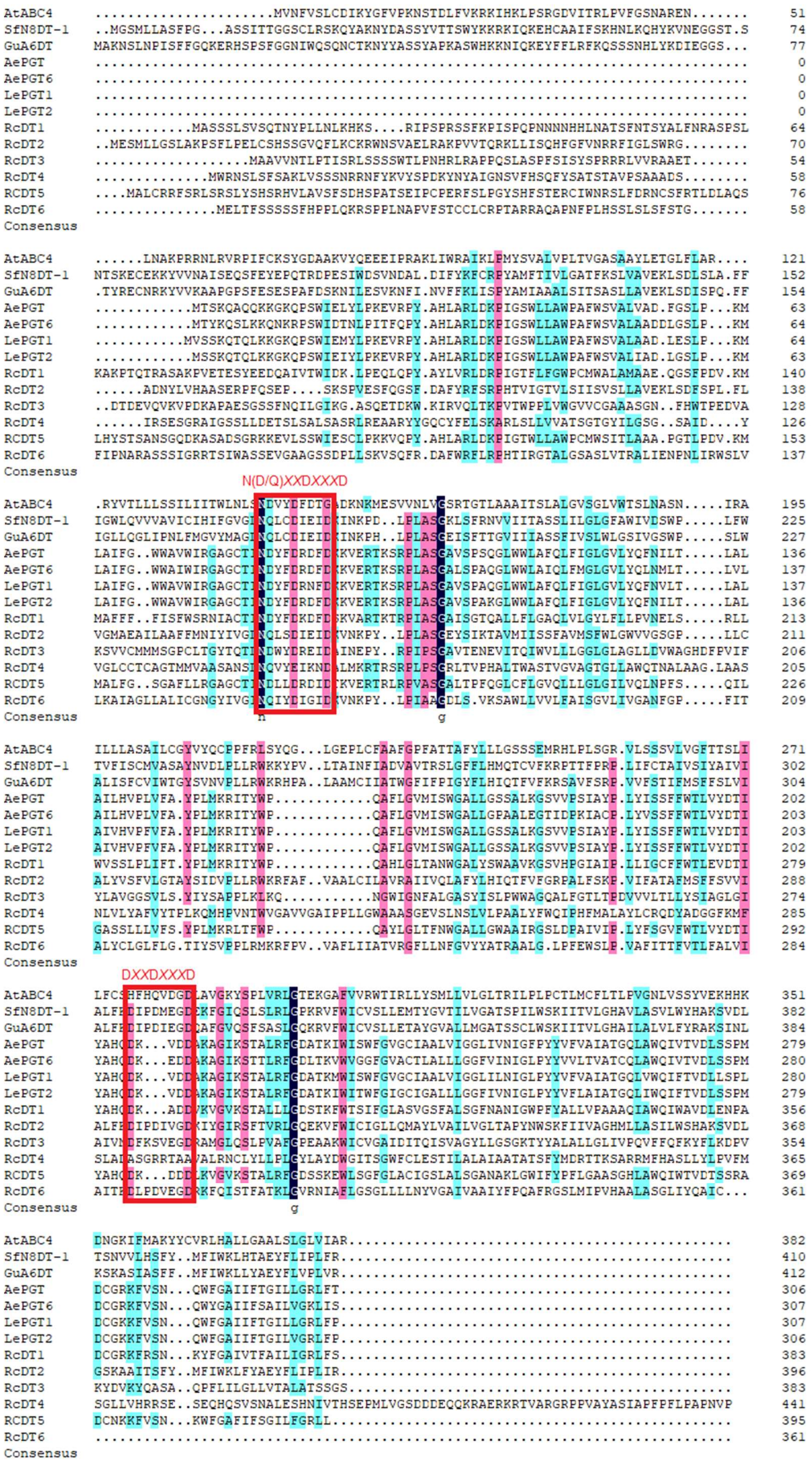
N(D/Q)XXDXXXD
DXXDXXXD

**Figure S1**. Alignment of the encoded polypeptides of the candidate genes. Amino acid sequences of a reported MenA homolog from *A. thaliana* (AtABC4, NM_001124046), a naringenin prenyltransferase from *Sophora flavescens* (SfN8DT-1, AB325579), a flavonoid prenyltransferase from *Glycyrrhiza uralensis*, and four 4-hydroxybenzoic acid geranyltransferases [LePGT1 (*L. erythrorhizon*, AB055078); LePGT2 (*L. erythrorhizon*, AB055079); AePGT (*A. euchroma*, DQ397513); AePGT6 (*A. euchroma*, KT991524)] are used for comparison with candidate genes. The conserved regions N(Q/D)*XX*D*XXX*D and D*XX*D*XXX*D are highlighted with red frames.

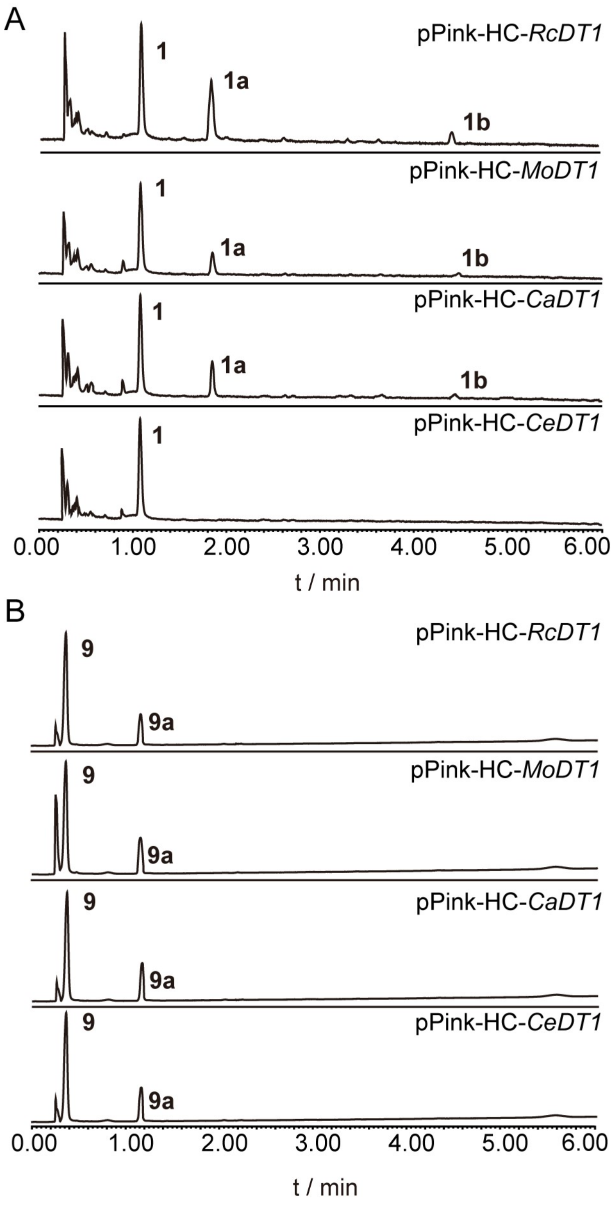


**Figure S2.** Comparison of the prenylation activities of recombinant RcDT1 and its rubiaceous homologs. A, UPLC-UV chromatograms of extracts from the prenylation reaction of 1,4-dihydroxy-2-naphthoic acid (**1**) catalyzed by recombinant RcDT1, MoDT1, CaDT1, and CeDT1. B, UPLC-UV chromatograms of extracts from the prenylation reaction of 4-hydroxybenzoic acid (**9**) catalyzed by recombinant RcDT1, MoDT1, CaDT1, and CeDT1. The UV detector was set at 254 nm.

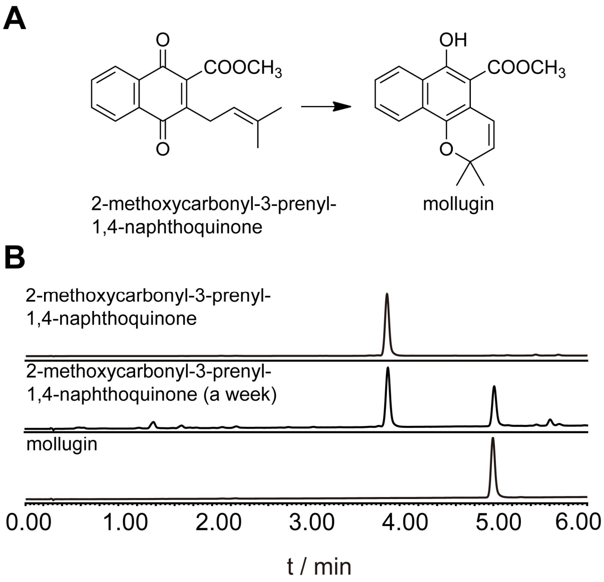


**Figure S3.** Spontaneous cyclization of 2-methoxycarbonyl-3-prenyl-1,4-naphthoquinone. A, Spontaneous cyclization reaction diagram of 2-methoxycarbonyl-3-prenyl-1,4-naphthoquinone. B, UPLC-UV chromatograms of authentic-standard 2-methoxycarbonyl-3-prenyl-1,4-naphthoquinone, 2-methoxycarbonyl-3-prenyl-1,4-naphthoquinone solution stored at 4 °C for one week, and authentic-standard mollugin. The UV detector was set at 254 nm.

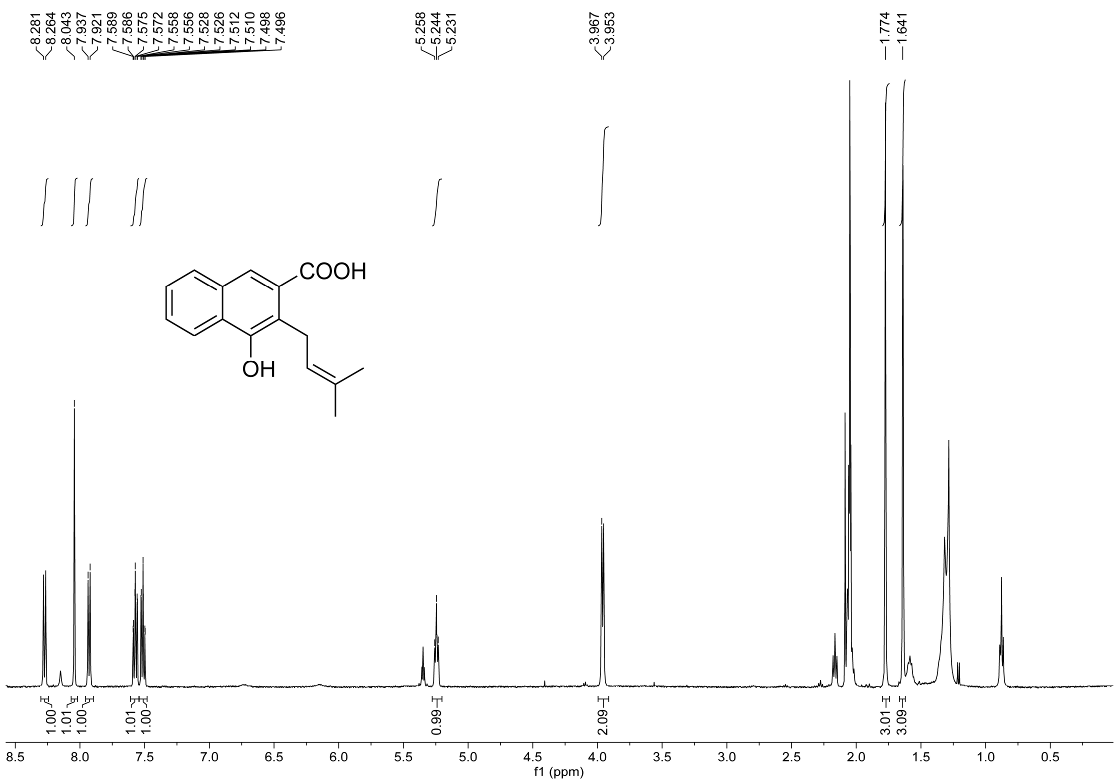


**Figure S4.** $^{1}H$ NMR spectrum of 3-prenyl-4-hydroxy-2-naphthoic acid (**2a**) in acetone-$d_6$ (500 MHz)

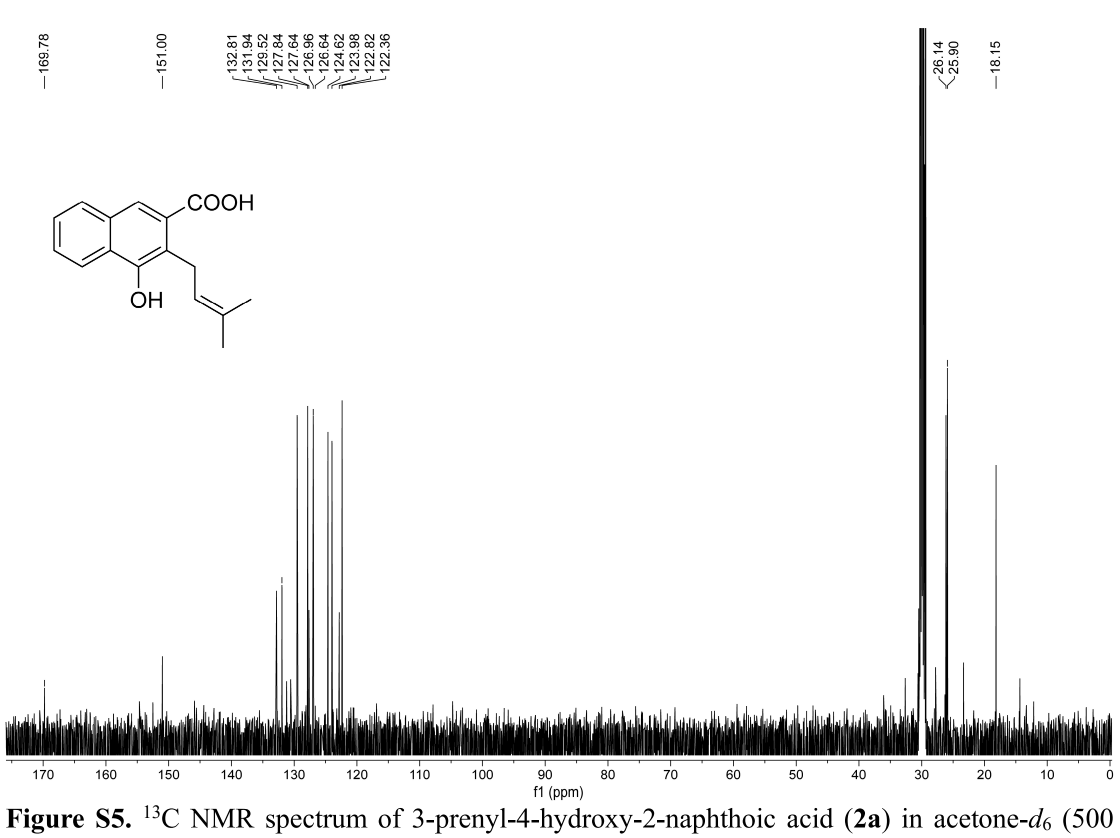


**Figure S5.** $^{13}C$ NMR spectrum of 3-prenyl-4-hydroxy-2-naphthoic acid (**2a**) in acetone-$d_6$ (500 MHz)

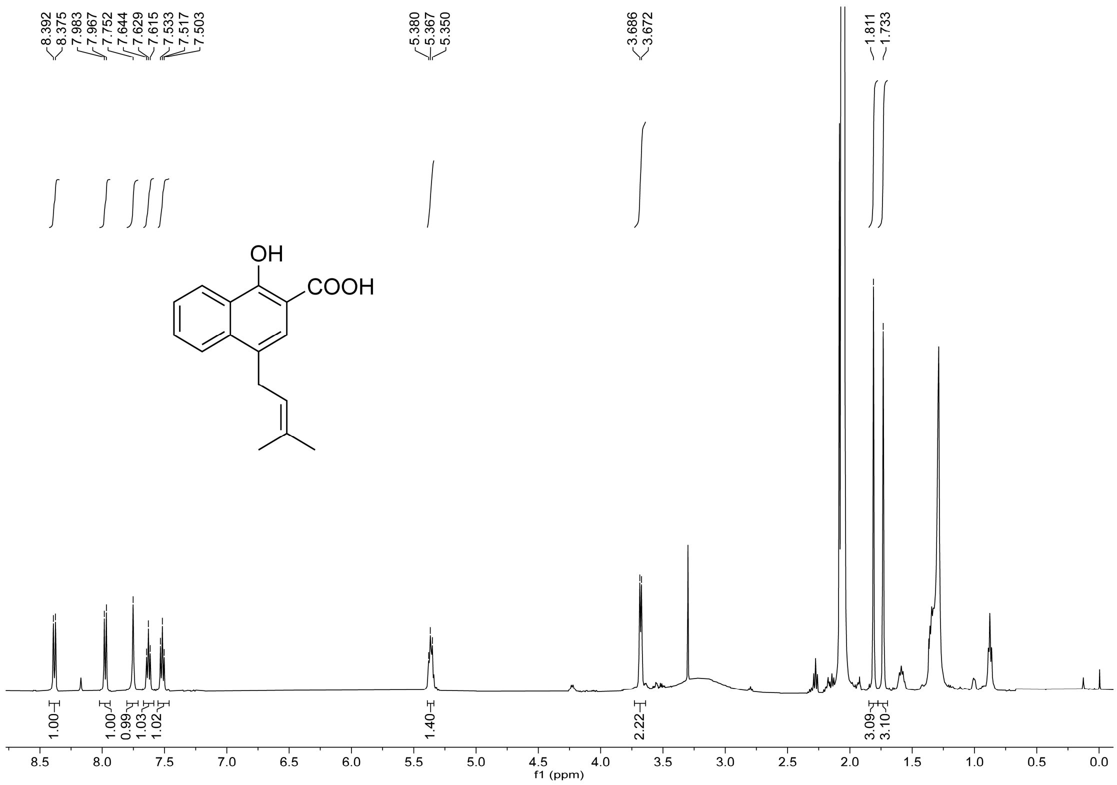


**Figure S6.** $^{1}H$ NMR spectrum of 4-prenyl-1-hydroxy-2-naphthoic acid (**3a**) in acetone-$d_6$ (500 MHz)

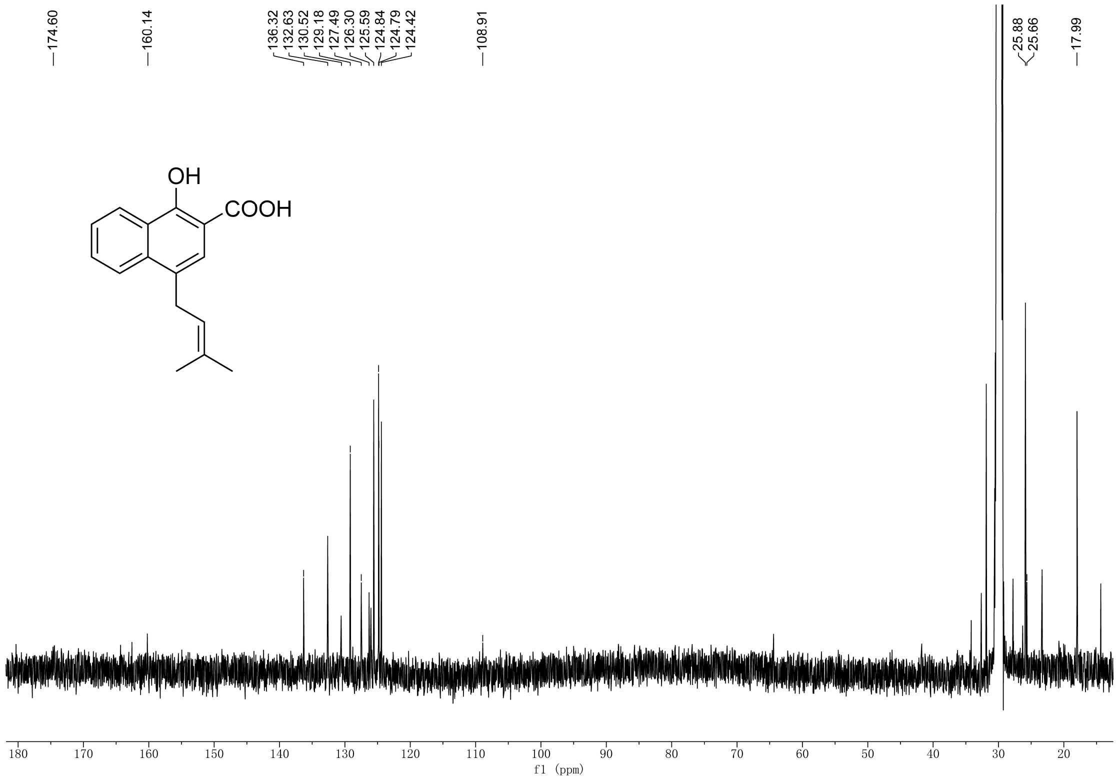


**Figure S7.** $^{13}C$ NMR spectrum of 4-prenyl-1-hydroxy-2-naphthoic acid (**3a**) in acetone-$d_6$ (500 MHz)

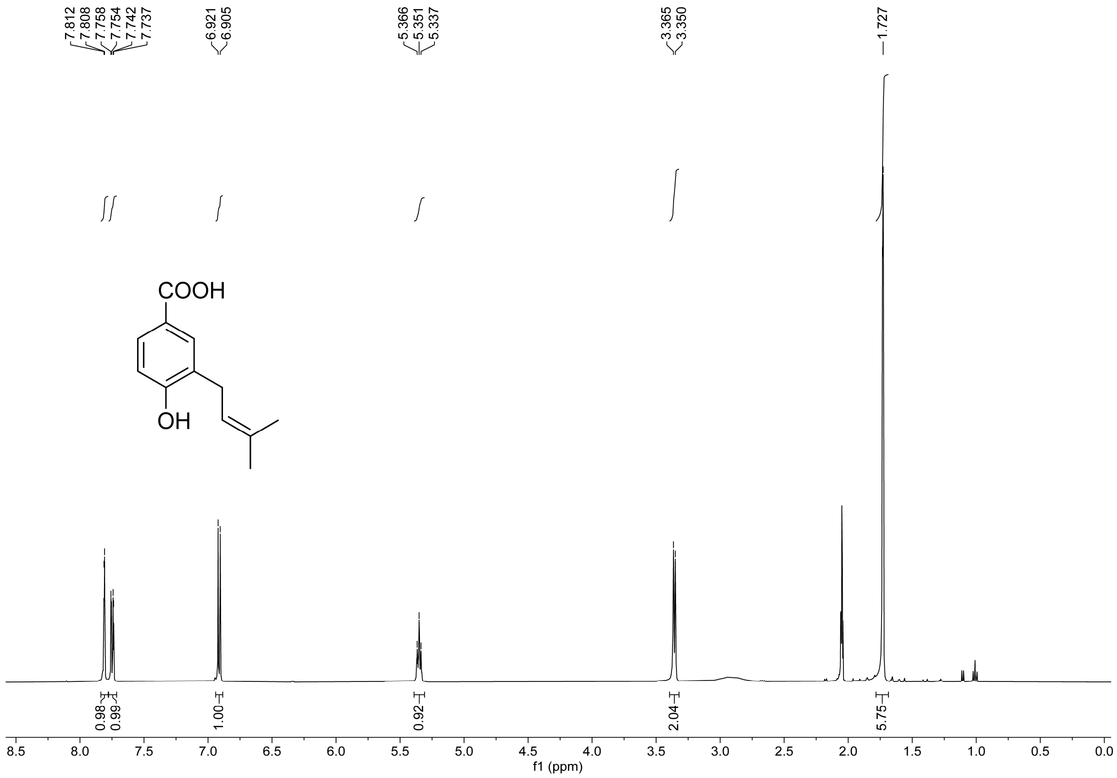


**Figure S8.** $^{1}H$ NMR spectrum of 3-prenyl-4-hydroxybenzoic acid (**9a**) in acetone-$d_6$ (500 MHz)

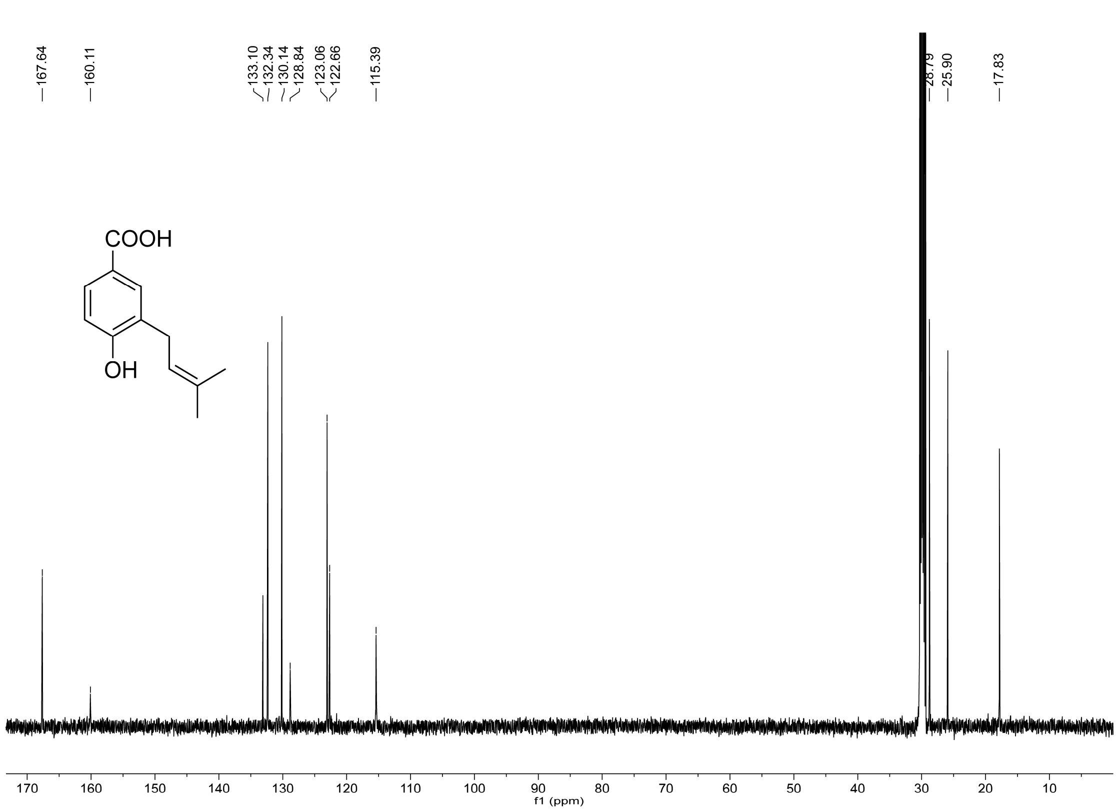


**Figure S9.** $^{13C}$ NMR spectrum of 3-prenyl-4-hydroxybenzoic acid (**9a**) in acetone-$d_6$ (500 MHz)

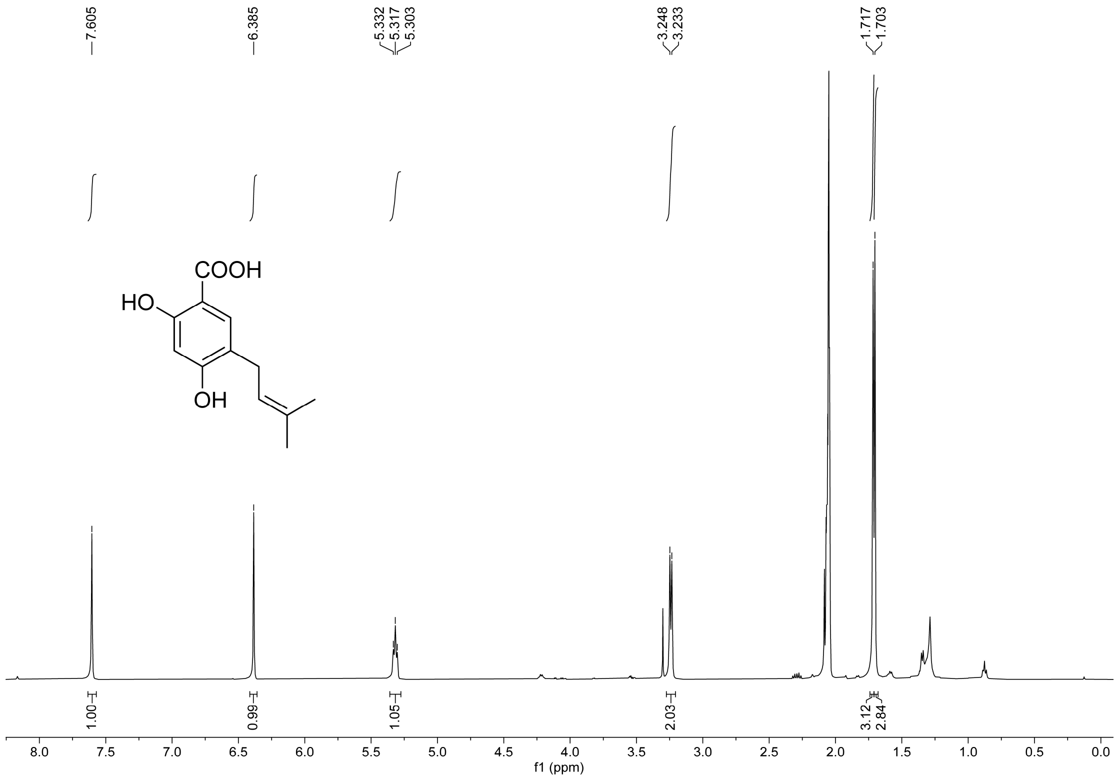


**Figure S10.** $^{1}H$ NMR spectrum of 5-prenyl-4-hydroxysalicylic acid (**11a**) in acetone-$d_6$ (500 MHz)

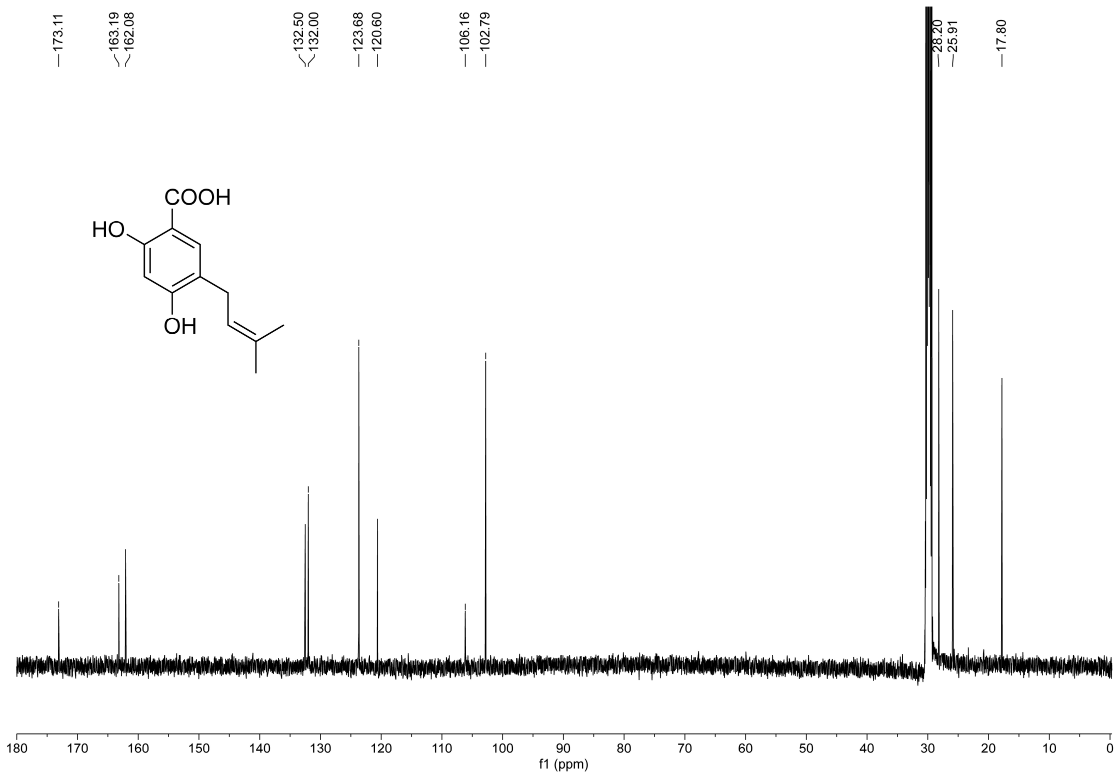


**Figure S11.** $^{13}C$ NMR spectrum of 5-prenyl-4-hydroxysalicylic acid (**11a**) in acetone-$d_6$ (500 MHz)

**Table S1.** The amino acid residues identities between RcDT1 and its rubiaceous homologs. 4 and 2 homologs with identities greater than 60% to RcDT1 are acquired from *C. arabica* and *C. eugenioides*, respectively. The homologs investigated in follow-up experiments are highlighted in red. The other genes are excluded due to their high similarity to the selected homologs.

| | Identities (%) | | | | | | |
|---|---|---|---|---|---|---|---|
| | *C. arabica* | | | | *C. eugenioides* | | *M. officinalis* |
| | CaDT3 (XP_027124881.1) | CaDT1 (XP_027122077.1) | CaDT2 (XP_027122550.1) | CaDT4 (XP_027124411.1) | CeDT1 (XP_027169671.1) | CeDT2 (XP_027169674.1) | MoDT1 |
| **RcDT1** | 64.78 | 66.67 | 61.26 | 60.10 | 61.15 | 60.10 | 73.79 |
| CaDT3 | — | 89.56 | 77.00 | 74.75 | 73.18 | 75.00 | |
| CaDT1 | | — | 76.23 | 78.82 | 72.39 | 79.13 | |
| CaDT2 | | | — | 72.19 | 69.57 | 72.19 | |
| CaDT4 | | | | — | 94.21 | 98.99 | |
| CeDT1 | | | | | — | 94.71 | |

**Table S2.** Primers used in this study.

| Applications | Name | sequence（5’-3’） |
|---|---|---|
| Full length cloning | RcDT1-F1 | ATGGCTTCCTCATCTCTCTCA |
| | RcDT1-R1 | CTAAGAAAAGAGCCTTCCAA |
| | RcDT2-F1 | ATGGAGTCTATGCTCCTGGG |
| | RcDT2-R1 | CTACCTTATAAGCGGTATGAGGA |
| | RcDT3-F1 | ATGGCCGCCGTCGTGAACAC |
| | RcDT3-R1 | CTAAGATCCGCTGCTAGTAG |
| | RcDT4-F1 | ATGTGGAGAAACTCACTGAGT |
| | RcDT4-R1 | TCAGTATGGTACATTCGGAGC |
| | RcDT5-F1 | ATGGCGTTATGCCGCCGTTT |
| | RcDT5-R1 | CTATAACAGGCGCCCAAATAG |
| | RcDT6-F1 | ATGGAGCTCACATTCTCGTC |
| | RcDT6-R1 | TCAGCATATTGCCTGGTAAATTAG |
| Subcloning | RcDT1-F24 | ATTCGAAACGGAATTCAAAAATGCCCAGCCCCAGATCCTC |
| | RcDT1-F48 | ATTCGAAACGGAATTCAAAAATGTCAGTTTCAACACCTCTTATGCGCT |
| | RcDT1-F72 | ATTCGAAACGGAATTCAAAAATGGCGTCGGCTAAGCCAGT |
| | RcDT1-R2 | TTAAATGGCCGGCCGGTACCCTAAGAAAAGAGCCTTCCAA |
| | RcDT2-F2 | ATTCGAAACGGAATTCAAAAATGGTGCAGTTTTTGAAATGCAAGA |
| | RcDT2-R2 | TTAAATGGCCGGCCGGTACCCTACCTTATAAGCGGTATGAGGA |
| | RcDT3-F2 | ATTCGAAACGGAATTCAAAAATGCGCCTTCGCGCTCCTCCACA |
| | RcDT3-R2 | TTAAATGGCCGGCCGGTACCCTAAGATCCGCTGCTAGTAG |
| | RcDT4-F2 | ATTCGAAACGGAATTCAAAAATGTACTCTCCTGATAAGTACAATTATGC |
| | RcDT4-R2 | TTAAATGGCCGGCCGGTACCTCAGTATGGTACATTCGGAGC |
| | RcDT5-F2 | ATTCGAAACGGAATTCAAAAATGTCTTTTTCCGACCACTCTCC |
| | RcDT5-R2 | TTAAATGGCCGGCCGGTACCCTATAACAGGCGCCCAAATAG |
| | RcDT6-F2 | ATTCGAAACGGAATTCAAAAATGCCAGTCTTCTCCACCTGTTG |
| | RcDT6-R2 | TTAAATGGCCGGCCGGTACCTCAGCATATTGCCTGGTAAATTAG |
| qRT-PCR | RcDT1-F1q | TTGATACCATTTATGCCCACCA |
| | RcDT1-R1q | CCAACCGAAGCAAGTCCAAA |
| | Actin-Fq | GATTGAGCACGGTATTGTTAG |

| | |
|---|---|
| Actin-Rq | ACACCATCACCAGAATCCAAC |

**Table S3.** The sequences used in this study.

| Name | sequence（5’-3’） |
|---|---|
| RcDT1 | ATGGCTTCCTCATCTCTCTCAGTCTCCCAAACCAACTATCCCCTCTTA<br>AACCTGAAGCACAAATCTCGTATTCCCAGCCCCAGATCCTCCTTCAA<br>ACCAATCTCACCTCAACCAAACAACAACAACCACCACCTTAACGCTA<br>CTAGTTTCAACACCTCTTATGCGCTTTTCAACCGTGCCTCCCCTTCTCT<br>GAAGGCTAAGCCGACGCAGACTCGTGCGTCGGCTAAGCCAGTGGAG<br>ACGGAGAGCTACGAGGAGGATCAGGCGATCGTGACGTGGATCGATA<br>AGCTGCCGGAGCAGCTTCAGCCGTACGCGTACCTTGTCCGGCTGGAC<br>CGGCCCATCGGGACGTTCCTGTTCGGGTGGCCGTGTATGTGGGCCCT<br>GGCCATGGCGGCGGAGCAAGGGAGCTTCCCCGACGTGAAGATGATG<br>GCCTTCTTCTTCTTCATCTCATTCTGGTCCAGGAATATTGCTTGCACCA<br>TTAATGACTATTTTGACAAGGACTTCGATTCTAAGGTGGCGAGAACA<br>AAAACAAGGCCAATAGCATCCGGAGCTATTTCTGGAACTCAAGCTCT<br>GCTTTTCCTTGGGGCTCAATTGGTACTCGGCTATCTGTTTCTTCTCCCG<br>GTCAACGAACTCAGCCGCCTCTTATGGGTTTCTTCGCTGCCTTTGATC<br>TTTACATACCCGCTGATGAAGAGAATAACATACTGGCCTCAAGCCCAC<br>CTGGGCCTAACAGCCAACTGGGGAGCTCTCTATTCTTGGGCCGCTGT<br>GAAAGGAAGTGTTCACCCTGGCATTGCCATCCCTCTCCTTATTGGTTG<br>TTTCTTCTGGACTCTTGAAGTTGATACCATTTATGCCCACCAGGACAA<br>AGCCGATGATGTGAAAGTAGGAGTGAAATCAACAGCATTGTTGCTTG<br>GAGATTCAACTAAGTTCTGGACAAGTATTTTTGGACTTGCTTCGGTTG<br>GAAGTTTTGCGCTAAGCGGTTTTAATGCTAATATTGGATGGCCATTCTA<br>TGCACTTTTGGTACCTGCGGCCGCTCAAATAGCGTGGCAGATATGGG<br>CTGTTGACCTAGAAAATCCAGCTGATTGTGGCAGAAAATTCCGATCC<br>AACAAATATTTCGGAGCAATTGTCACATTTGCCATTCTAATTGGAAGG<br>CTCTTTTCTTAG |
| RcDT2 | ATGGAGTCTATGCTCCTGGGTTCCTTGGCAAAACCTTCTTTTTTACCT<br>GAACTCTGTTCACATTCTTCAGGGGTGCAGTTTTTGAAATGCAAGAG<br>ATGGAACAGTGTGGCGGAACTACGCGCAAAGCCTGTGGTCACTCAG<br>AGGAAGCTTCTGATTAGCCAGCATTTTGGTTTTGTTAATAGGAGATTT<br>ATTGGGCTTTCTTGGAGAGGAGCTGATAATTATCTGGTGCATGCGGCC<br>TCTGAACGCCCTTTTCAATCTGAGCCTTCAAAGAGCCCCGTGGAATC<br>ATTTCAAGGGAGTTTTGATGCTTTCTATCGGTTTTCGCGGCCCCATAC<br>AGTCATTGGAACTGTGCTGAGCATAATTTCAGTTTCTCTTCTAGCAGT<br>GGAGAAGCTCTCGGATTTTTCACCACTGTTTCTTGTTGGGATGGCTGA<br>GGCCATTCTTGCAGCCTTCTTCATGAACATATACATTGTTGGTTTGAAT<br>CAGTTGTCAGACATAGAAATCGACAAGGTTAACAAGCCTTACCTTCC<br>ATTGGCATCAGGAGAATATTCAATCAAGACGGCGGTGATGATTATTTC<br>ATCTTTTGCAGTAATGAGTTTTTGGCTCGGATGGGTTGTTGGCTCTGG<br>CCCATTACTTTGTGCTCTTTATGTCAGTTTTGTGCTTGGGACAGCATAT<br>TCAATCGACGTACCCTTGTTGAGATGGAAGAGGTTTGCGTTTGTTGC<br>CGCACTCTGTATCTTGGCTGTGCGAGCAATAATTGTACAGTTAGCATT<br>TTATTTGCACATACAGACTTTTGTGTTTGGACGGCCAGCTCTCTTTTC<br>AAAGCCTGTAATTTTTGCAACAGCATTTATGTCCTTCTTCTCAGTTGTA<br>ATAGCACTTTTTAAGGATATTCCTGATATTGTTGGAGACAAAATTTATG<br>GCATCCGATCTTTTACGGTCAGGTTGGGTCAAGAGAAGGTCTTCTGG<br>ATTTGCATTGGACTTCTTCAAATGGCTTACCTTGTTGCCATTTTAGTCG<br>GTTTGACAGCACCCTATAACTGGAGCAAATTCATAATAGTTGCTGGTC<br>ACATGCTTTTGGCATCGATACTTTGGAGTCATGCCAAATCTGTTGATC<br>TAGGAAGCAAGGCAGCAATAACATCCTTTTACATGTTTATATGGAAGC<br>TTTTTTATGCCGAGTACTTCCTCATACCGCTTATAAGGTAG |

| | |
|---|---|
| RcDT3 | ATGGCCGCCGTCGTGAACACACTTCCCACTATCAGTAGATTATCCAGT<br>TCCAGCTGGACACTACCCAACCACCGCCTTCGCGCTCCTCCACAGTC<br>ACTCGCTTCACCATTTTCAATCTCTTATTCTCCCCGAAGAAGGTTAGT<br>GGTGAGAGCTGCAGAGACTGATACTGATGAAGTACAAGTTAAGGTAC<br>CGGATAAGGCACCTGCGGAGAGTGGTTCCAGCTTTAACCAGATTCTC<br>GGGATCAAAGGCGCCAGCCAAGAAACAGATAAATGGAAGATTCGGG<br>TTCAACTTACGAAGCCGGTTACTTGGCCTCCTCTTGTGTGGGGTGTTG<br>TATGTGGCGCTGCAGCTTCTGGGAACTTCCACTGGACACCGGAGGAT<br>GTAGCCAAATCAGTGGTCTGCATGATGATGTCAGGCCCCTTGTCTAACT<br>GGATATACTCAGACACTAAATGACTGGTATGATAGAGAGATTGATGCG<br>ATCAATGAGCCTTATCGGCCAATCCCTTCTGGAGCAGTAACTGAGAA<br>CGAGGTGATCACCCAAATATGGGTCCTGCTTCTTGGTGGCCTAGGTTT<br>GGCTGGGCTTTTGGATGTTTGGGCAGGGCATGATTTCCCGGTGATATT<br>TTACCTTGCTGTCGGGGGCTCTGTGCTATCCTATATCTATTCAGCTCCT<br>CCTCTAAAGCTTAAACAGAATGGATGGATTGGCAATTTCGCTCTTGGA<br>GCAAGTTATATCAGTTTGCCATGGTGGGCGGGTCAGGCTCTCTTTGGC<br>ACCCTTACACCCGACGTAGTCGTTCTCACTCTCTTGTATAGTATCGCC<br>GGGCTTGGCATTGCCATTGTGAATGACTTCAAAAGTGTTGAAGGAGA<br>CAGAGCTATGGGGCTTCAGTCCCTTCCAGTTGCTTTTGGACCTGAAG<br>CTGCTAAATGGATCTGTGTCGGAGCCATCGACATAACCCAGATATCAG<br>TTGCAGGCTATCTTCTTGGTAGCGGTAAAACTTACTACGCACTAGCCC<br>TACTTGGTTTAATTGTACCCCAAGTATTTTTCCAGTTCAAGTACTTCCT<br>CAAGGACCCTGTGAAATACGATGTCAAGTATCAGGCTAGTGCTCAGC<br>CGTTTCTTATTCTTGGCCTTCTCGTTACCGCTTTAGCTACTAGCAGCGG<br>ATCTTAG |
| RcDT4 | ATGTGGAGAAACTCACTGAGTTTCTCAGCCAAACTTGTTTCTTCCTCC<br>AACCGTAGGAACTTTTACAAGGTTTACTCTCCTGATAAGTACAATTAT<br>GCCATTGGAAACTCGGTTTTCCACTCTCAATTTTATTCGGCGACTTCA<br>ACAGCAGTACCTTCTGCTGCTGCTGACTCCATCAGATCAGAGTCCGG<br>TCGAGCTATCGGATCCTCTTTACTCGATGAGACGTCGTTGTCTGCTTT<br>GTCGGCCAGTAGGTTGAGAGAAGCTGCCAGATATTACGGTCAATGTT<br>ATTTTGAACTCTCCAAAGCTCGACTTAGTTTGTTAGTCGTTGCTACTT<br>CGGGTACTGGATATATCCTTGGTAGTGGCAGTGCCATAGATTACGTTG<br>GACTGTGTTGCACGTGTGCTGGTACGATGATGGTTGCAGCATCTGCA<br>AACAGCTTAAATCAGGTTTATGAAATAAAAAATGATGCCCTTATGAAG<br>AGAACCAGGAGCAGACCATTACCTTCTGGGCGTCTCACTGTTCCTCA<br>TGCACTCACCTGGGCTTCTACGGTTGGAGTTGCTGGGACTGGTTTAC<br>TTGCATGGCAGACTAATGCGTTGGCTGCTGGCCTTGCTGCTTCTAATC<br>TCGTTCTTTATGCATTTGTATACACTCCCCTGAAACAAATGCATCCAGT<br>TAATACTTGGGTGGGCGCTGTTGTTGGTGCTATTCCGCCTCTACTTGG<br>GTGGGCTGCTGCCTCCGGCGAAGTTTCACTAAATTCTCTTGTTCTCCC<br>TGCTGCTTTATATTTTTGGCAAATACCTCATTTCATGGCCCTAGCATAC<br>TTGTGTCGTCAAGACTATGCAGATGGAGGGTTTAAAATGTTCTCCCTT<br>GCTGATGCTTCTGGTCGGAGAACTGCTGCCGTTGCCTTGAGGAACTG<br>CTTGTATCTGCTTCCATTGGGCTATCTAGCCTACGACTGGGGAATAAC<br>ATCTGGCTGGTTCTGCCTCGAATCCACAATCCTCGCTCTGGCAATTGC<br>TGCCACCGCCACATCTTTCTACATGGATCGCACAACAAAGAGCGCGA<br>GGAGAATGTTCCACGCTAGTCTCCTGTACCTTCCTGTATTCATGTCCG<br>GACTTCTGGTTCATCGTCGATCAGAAAGCGAGCAGCACCAAAGTGTT<br>AGCAATGCCCTCGAGTCTCACAATATCGTGACACACTCAGAACCAAT<br>GCTTGTTGGGAGTGATGACGACGAACAGCAGAAAAGGGCCGAAAGA<br>AAAAGAACGGTTGCGCGTGGAAGACCTCCTGTGGCGTATGCATCCAT<br>CGCGCCTTTCCCTTTCTTACCAGCTCCGAATGTACCATACTGA |
| RcDT5 | ATGGCGTTATGCCGCCGTTTCTCCCGTCTATCCCGCTCGCTCTACTCCC<br>ACTCCCGCCATGTTTTAGCAGTTTCTTTTTCCGACCACTCTCCGGCTA<br>CAAGTGAAATTCCATGTCCGGAACGTTTTTCCCTACCTGGTTATTCAC<br>ATTTCAGTACAGAACGCTGCATTTGGAATAGGAGCCTATTTGACAGA |

| | |
|---|---|
| | AATTGTAGTTTCAGAACTCTAGACCTAGCTCAATCACTTCATTATTCA<br>ACTTCAGCGAATTCGGGACAGGACAAAGCTAGTGCAGATAGTGGTAG<br>AAAGAAGGAAGTTTTGTCGTCTTGGATTGAATCGTGCTTACCGAAAA<br>AGGTTCAGCCATACGCACACTTAGCGAGGCTGGATAAGCCGATAGGG<br>ACTTGGTTGTTAGCCTGGCCTTGTATGTGGTCTATTACTTTGGCAGCT<br>GCTCCAGGAACCCTCCCTGATGTGAAAATGATGGCACTTTTTGGTTCC<br>GGGGCATTTTTATTGCGAGGTGCTGGTTGTACCATTAATGATCTGCTT<br>GATAGGGATATCGATACTAAGGTAGAGCGGACTAGGTTGAGGCCAGT<br>TGCCAGTGGTGCGTTGACACCTTTTCAGGGGCTTTGTTTTCTCGGCGT<br>GCAGTTGCTTTTGGGACTTGGGATTCTTGTTCAGCTGAATCCCTTCAG<br>TCAAATCTTAGGTGCGTCGTCTCTGTTGTTGGTATTCTCATATCCTTTA<br>ATGAAACGGCTGACATTTTGGCCTCAAGCCTATCTCGGTTTAACTTTT<br>AACTGGGGAGCCTTGCTTGGCTGGGCTGCTATTAGAGGGAGTCTTGA<br>TCCTGCGATTGTGATCCCTCTTTATTTTTCTGGTGTGTTTTGGACACTT<br>GTGTACGATACCATCTATGCACACCAGGATAAAGACGATGACCTGAA<br>AGTGGGCGTCAAATCAACAGCACTGAGGTTCGGGGATTCAAGCAAA<br>GAATGGCTTTCTGGATTTGGACTCGCCTGCATAGGTAGTCTTGCCCTC<br>AGCGGGGCAAATGCCAAACTGGGATGGATCTTTTATCCATTTTTAGGA<br>GCTGCATCTGGTCACTTAGCTTGGCAGATATGGACGGTTGATACATCG<br>TCCCGGGCTGATTGCAATAAGAAATTCGTCTCCAATAAATGGTTTGGT<br>GCGTTTATCTTCAGTGGTATCCTATTTGGGCGCCTGTTATAG |
| RcDT6 | ATGGAGCTCACATTCTCGTCTTCTTCTTCATTCCATCCTCCTTTGCAAA<br>AGCGCTCCCCTCCATTGAATGCTCCAGTCTTCTCCACCTGTTGCCTCT<br>GTAGGCCCACAGCAAGAAGAGCCCAAGCTCCGAACTTTCCACTCCAT<br>TCTTCCCTTTCCCTGTCCTTCTCAACTGGGTTTATCCCCAATGCCAGA<br>GCATCATCCTCCATTGGAAGGCGCACTTCCATCTGGGCTTCCTCCGAA<br>GTTGGTGCTGCTGGATCCTCTGATCCCTTGTTAAGCAAAGTATCGCAG<br>TTCAGAGATGCCTTCTGGAGGTTTCTGAGGCCTCACACTATCCGTGG<br>CACAGCATTAGGATCAGCCTCGTTAGTGACGCGCGCTCTGATTGAAA<br>ACCCAAATCTGATAAGATGGTCACTTGTGTTAAAGGCTATAGCCGGTC<br>TTCTTGCTCTTATATGCGGGAATGGTTACATAGTGGGAATCAATCAGAT<br>ATATGACATTGGTATTGATAAGGTAAACAAACCTTATTTGCCTATTGCT<br>GCAGGAGATCTTTCTGTCAAATCTGCTTGGCTTTTGGTGGTGCTCTTC<br>GCCATCTCTGGTGTTTTGATTGTTGGAGCAAACTTTGGTCCATTTATC<br>ACGGCCCTTTATTGCCTGGGTCTCTTTCTAGGGACTATTTATTCTGTTC<br>CGCCACTTCGGATGAAGAGATTTCCAGTTGTGGCATTTCTTATAATTG<br>CCACGGTTCGTGGTTTCCTTCTTAACTTTGGGGTTTATTATGCTACAAG<br>AGCTGCTCTTGGGCTCCCATTTGAGTGGAGCTTACCAGTGGCTTTCAT<br>CACGACATTTGTAACATTGTTTGCACTGGTCATTGCCATAACGAAGGA<br>TCTTCCAGACGTGGAGGGTGATCGCAAGTTTCAGATTTCTACATTTGC<br>CACAAAGCTCGGCGTGAGAAATATAGCATTTCTTGGTTCCGGGCTATT<br>ACTTCTAAACTACGTTGGTGCCATTGTAGCAGCTATTTATTTTCCTCAG<br>GCATTCAGGGGAAGCTTGATGATTCCCGTACATGCAGCTTTAGCATCG<br>GGTCTAATTTACCAGGCAATATGCTGA |
| CaDT1 | ATGGCTGTATTGGTGATCGCTTCCTTCCTGATAGCCTATTGTTGTGGGA<br>GGTACTCGAGGGATGGCAGCTCACTTGCTGGGAGCGAGAGTGGGAA<br>GAGGAGGAGTAGTTATTCCGGAGTTAAGAGGAGGGTTTATGCCAACA<br>ATGGAAGTTTGATGGAAATTACAAGAGGCTTCAACTTAGCTAATCAA<br>GCTCCACAAGTTTCAGCCTCAGATAAGACAATAGAGAAAGCTGAAG<br>AAGAGAAAGCTGTCACTTCTTCTTGGATTGAAGCAGTTTTTCCTGAA<br>ACAGCCCGGCCTTATGCCTACCTTGTCCGCTTGGACAACCCGACCGG<br>GACGTTTTTATTTGCATGGCCATGCTTGTGGTCGCTTGCATTCACTGC<br>GAACCCCGGCAGCCTTCCTGATATGAAGATGTTGGCATTCTTTTTCCT<br>GGTGTCTTTTATGTCAAGAAATATTGCATGCACCATTAACGATTATTTT<br>GACAAGGATTTTGATGCACAGGTTGAGCGAACCAAAGGAAGGCCGC<br>TTGCCTCTGGGGCAATAACGGGATTTCAAGCACTTTGCTTCCTTGGA<br>ATTCAAGTGTTACTCGGATATGGAATTTTTCTCCCAATAAATGAACTA |

| | |
|---|---|
| | AGCCGTATTTTATGGGTTTCATCCTTGCCGTTGATCTTCACTTACCCGC<br>TCATGAAGAGAATTACTTATTGGCCTCAAGCCCATCTTGGTTTAACTG<br>CAAATTGGGGAGCTTTGTACTCATGGGCTGCTGTTAAAGGAAGTCTT<br>GATCCTGCTATTGTCTTCCCGGTCCTCGTTGCTTGCTTCTTCTGGACA<br>CTGGAGGTGGATACAATATATGCACATCAGGATAAAGAAGATGATGTG<br>AAAGTGGGTGTTAAATCTACAGCTTTGCTATTAGGTGATTCAACAAAA<br>TTGTGGACTACTGGTTTTGGAGTTGCATCCATTGCTAGTCTTGCTCTA<br>GCTGGATTTAACGCTCACATTGGATGGCCATTTTTTGTATTGCTAGCG<br>GCTGCATCAGGTCAAATAGCTTGGCAGATTTGGGATGTTGACTTATCA<br>AACCCAGCAGATTGTTTCAGAAAATTTGCATCAAATAGATATTTTGGT<br>GCTATTGTCTTTAGTGCAATTCTGTTTGGAAGACTCTTGTCATAA |
| CaDT2 | ATGGCATCCTCTCATAGTTTCTCTGGGATTTTTCCCATCTCCAACTTCA<br>ACCATCAACCTCATCCCTCCATAGCTCTCTCCAAATCCTCCATCAAAC<br>CAGCCAAATCAATCTCCAAAATCAACCCTTCTAGTGTGGAAAACTCA<br>GCCACATTCAACCATTCTTCGATAAAGGATACCAGATTTCTTGGTAGA<br>GGCTGTAACCTAACATATCAAGCTCCACGATTATCAGCATCATCAAAG<br>TCATTAGAGCAGAGTGAAGATGAAAAACCTGCTTCTTCTTCTTGGATT<br>GAATCGGTTTTTCCTGAAAAAGCTCGGCCTTATGCCTACCTAGTCCGG<br>TTGGACAAGCCGATCGGCACGTTGTTAATTGCATGGCCATGTATGTGG<br>TCACTAGCATTCGCGGCGAACCCCGGCAGCCTTCCTGATGTGAAGAT<br>GTTGGCATTCTTTTTCTTGGTGTCTTTCTTGACGAGAAACATTGGATG<br>CACCATTAACGATTACCTTGATAAGGATTTCGATGCACAGGTGGAGAG<br>AACCAAAGGAAGGCCTCTGGCCTCTGGCTCAATCACCGGATTTCAAG<br>CCCTTCGTTTCCTTGGCATCCAATTGCTAATTGGTTATGGAGTTCTTCT<br>CCCAGTAAACGAACTAAGCCGTCTACTATGGGCTTCATCCTTACCGTT<br>GATCTTCATTTACCCGCTTATGAAGAGAATAACATACTGGCCTCAAGC<br>CCTATTGGGACTGGCTATGAACTGGGGAGTATTATGCTCCTGGGCTGC<br>TGTGAAAGGAAGTCTTGATCCTACTATTGTTTTCCCACTTCTTGTTGG<br>ATGCTTCTTTTGGACTCTAGAGTATGATACAATATATGCACATCAGGAT<br>AAAAAGGATGACGTGAAAGCGGGTGTCAAATCTACAGCTTTGCTATT<br>GGGCGAATCAACAAAATTGTGGTCTGCTAGTTTTGGAGCGGCATCCA<br>TTGCTTGTTTTGCTCTAGCCGGATTGAATGCTCATATTGGATGGCCATT<br>TTATGCATTGCTAGTAGCTGCTTCCGGTCACATAGCTTGGCAAATTTG<br>GGATGTTGACCTATCAAACCCGGCAGACAGTTTCAGAAAATTTGCAT<br>CAAATAAATGGTTTGGTGGAATTGTCTTTAGTGCAATCCTATTCGGAA<br>GATTCCTTTCCTAG |
| CeDT1 | ATGGCCCAAGCTTCCAAGCTTATGGCTTCCTCTCACAGTAGTGTTCAT<br>TTCATCTCCAGCTTGAACCATCAACGTCGCCCCTCCTTAGCTGATCTC<br>CCCAAATCCTTCCTGAAACCAGTCAAATCAAACTCCAAAATCAGCCC<br>TTGCACAATACTGGAAAATAGCTCAGCCAGCTTCCGCCGTTCTTCGGT<br>TGAACATATCAAATTCCTTGGCAGAGGCTGCTGCCACTTACCTATCCG<br>AGATCCACATTTTTCAGCCTCAGCAAAGTCCTTAGAGCGAGGCGAAG<br>GAGGACAACCCGTTCCTCCTTGGATTGAAGCAATTTTTCCTGAACAA<br>GCTCGGCCTTACGCCCACCTAGTCCGCTTGGACAAGCCAGTTGGGAC<br>GTTTTTATTTGCATGGCCATGTATTTGGTCGCTTGCATTCACGGCGAAT<br>GCCGGCACGCTTCCTGATCTGAAGATGTTAGCATTTTTTTTCTTCGTC<br>TCTTTCATGTCAAGAAATATTGCATGCACCATCAACGATTACTTTGATA<br>AGGATTTCGATTCGCAGGTTGAGCGAACCAAGGGAAGGCCGCTTGC<br>CTCCGGCGCAATAGCGGGATTTCAAGCACTTCTTTTCCTTGCCATTCA<br>ATTGCTACTCGGTTATGGTGTTCTTTTCGCAGTAAACAAACTAAGCCG<br>TCTATTATGGGTTTCATCTTTGCCGTGGATCTTTACTTACCCGCTCATG<br>AAGAGAATAACATATTGGCCTCAAGCCTATCTGGGTCTGACGGTCAG<br>TTGGGGCGCAGCATACTCATGGGCTGCTGTTAAAGGAAGTCTTGAGC<br>CTGCTATTGTCTTCCCACTCGTCCTTGCCTTCTTCTTTTGGACTCTGGA<br>ATTTGATACAATATATGCACATCAGGACAAAGAAGATGATGTGAAGGT<br>GGGCATCAAGTCTACAGCTTTGCTATTTGGTGATTCAACAAAACTGTG<br>GATTTCTGGTTTTGCAGCGGCATCCATTGTTAGTCTTGCTCTAACCGG |

| | |
|---|---|
| | ATTTAATGCTAACATTGGATGGCCATTCTATGGATTGTTGGCTGCTGCT<br>TCTGGTCATCTTGCTTGGCAAATTTGGGATGTTGACTTATCAAATCCT<br>GCTGATTGCTCCAGAAAATTTGTATCAAATATATGGTTTGGTGCTATTG<br>TCTTTGGTGCGATCTTATGTGGAAGACTCTTCTCATAG |
| MoDT1 | ATGGCTTCCTCCTTCTCTGTCTCTCAAACTTCCATCTTGAACTTGAAG<br>TATGTCCCTCGTACTTCCAAACCTCTCCCAAAATCGTCCCTTAAACCA<br>TTCCAATCAAACACAAACATTAGCCCATATAATGGCATCAACATTCCT<br>CAAATCCCAACAAATTTCAAAAACTCTTCAATTCTTTTCAACAAGTCT<br>TCATCATCAATCGAAAATTCGAAAGTTCTTGCTAAACCTCCACAGTTC<br>TCAGCCTCAACAAAGACAGTGGAGGAAAATGTGGAGGAAGACCAAC<br>TGATGGTTTCTTGGATCGATTTGTTGCCCGAGAAAATTCAGCCATATG<br>CTTATTTGGTCCGGTTGGACAGGCCAATCGGAACATTTTTATTTGGAT<br>GGCCGTGTGTGTGGTCACTAGCACTCGCGGCGGAACCCGGCATGTTG<br>CCTGACTGGAAGATGTTGGCATACTTTTTCTTTGTGTCTTTCTGGTCTA<br>GAAATATTGCATGCACCATCAATGACTATTTTGACAAGGATTTTGATTC<br>AAAGGTTGAGCGAACAAAAAGGAGGCCGATTGCCTCTGGGGCTATT<br>TCAGGACTTCAAGCACTTATTTTCCTTGGTGTTCAAGTGGTGCTCGGT<br>TATGCCATTCTTCTCCCCGTCAACGAATTAAGCCGCCTATTATGGATTT<br>CTTCGTTGCCATTGCTCTTTACCTACCCGCTCATGAAGAGAATAACAT<br>ATTGGCCTCAAGCCCATCTTGGTCTGACTGCCAACTGGGGAGCGTTG<br>TATTCTTGGGCTGCTGTTAAAGGAGCTCTTCATCCTTGGATTGCCATC<br>CCACTTCTCATTGGTTGCTTCTTCTGGACTCTTGAGGTTGACACAATT<br>TACGCCCATCAGGATAAAGAAGATGATGCGAAAGTAGGAGTCAAATC<br>TACAGCATTGTTGCTTGGCGATTCGACAAAATTGTGGACCACTGTTTT<br>TGGAATCGCATCCATCGCTAGTTTTGCATTCAGTGGATTCAATGCTCAT<br>ATTGGATGGCCGTTTTATGCACTTCTGGTACCTGCTGCTGCTCAAATA<br>GCTTGGCAGATTTGGGCGGTGGACTTAGAAAACCCAGCAGATTGCGG<br>CAGAAAATTTCGATCCAACAAATATTTTGGAGGGATTGTCTTCTTCGC<br>CATTCTATTCGGAAGACTGTTCTCATAG |
